# Systemic immune response profiling with SYLARAS implicates a role for CD45R/B220^+^ CD8^+^ T cells in glioblastoma immunology

**DOI:** 10.1101/555854

**Authors:** Gregory J. Baker, Jeremy L. Muhlich, Sucheendra K. Palaniappan, Jodene K. Moore, Stephanie H. Davis, Sandro Santagata, Peter K. Sorger

## Abstract

Accurately profiling systemic immune responses to cancer initiation and progression is necessary for understanding tumor surveillance and, ultimately, improving therapy. Here, we describe the SYLARAS software tool (<u>SY</u>stemic <u>L</u>ymphoid <u>A</u>rchitecture <u>R</u>esponse <u>AS</u>sessment) and a data set collected with SYLARAS that describes the frequencies of immune cells in primary and secondary lymphoid organs and in the tumor microenvironment of mice engrafted with a standard syngeneic glioblastoma (GBM) model. The data resource involves profiles of 5 lymphoid tissues in 48 mice and shows that GBM causes wide-spread changes in the local and systemic immune architecture. We perform in-depth analysis of one significant tumor-induced change: depletion of a specialized subset of CD45R/B220^+^ CD8^+^ T cells from the circulation and their accumulation in the tumor mass. Immunoprofiling of tissue microarrays demonstrates the presence of similar cells in human GBM.

## INTRODUCTION

Glioblastoma (GBM) is an aggressive and incurable brain tumor characterized by high intrinsic and adaptive immunotherapeutic resistance^1^. Like many solid cancers, it dampens the effector function of tumor-resident immune cells by producing anti-inflammatory cytokines/catabolites^2,3,4,5,6^, lectins^7,8^, and immune checkpoint molecules^9,10^. Interest in using immunotherapy to treat GBM is driven by evidence of dramatic tumor regression in orthotopic immunocompetent murine models^11^ and encouraging but sporadic responses to immune checkpoint inhibitors (ICIs) in human patients^12,13,14,15^. However, the success of ICI therapy for GBM and other tumors of the central nervous system likely depends on a more complete description of immune cell interactions within and across lymphoid tissues in response to tumor growth, the cell and molecular repertoires necessary for efficacious ICI therapy, and biomarkers predictive of ICI response. In this paper we tackle the first of these challenges.

The immune system comprises a complex network of specialized cells that communicate with each other and traffic to distinct tissues to confer resistance to foreign and self-antigens. Key primary and secondary lymphoid tissues include the blood, bone marrow, lymph nodes, spleen, and thymus each of which plays complementary roles in the priming and maintenance of robust anti-tumor immunity. Despite this, cancer immunology has focused primarily on tumor-infiltrating immune cells and their behavior within the tumor microenvironment (TME). Recent results from animal models of cancer show that effective immunotherapy depends on the peripheral immune system^16^, although the effect of cancer on immunological events taking place across the peripheral immune system remains unclear. This is due in large part to a lack of effective tools for processing, analyzing and visualizing large sets of immuno-profiling data characterizing multiple lymphoid organs across time and disease status.

Here we describe SYLARAS (<u>SYs</u>temic <u>L</u>ymphoid <u>A</u>rchitecture <u>R</u>esponse <u>AS</u>sessment), a tool for studying systemic immune responses. SYLARAS combines multiplex immunophenotyping with software for transforming complex single-cell datasets into a visual compendium of time and tissue-dependent changes in immune cell frequencies and the relationships between these frequencies. We focus on perturbations imposed by GBM, but our approach is applicable to other cancers, infectious or autoimmune disease, vaccines, immunotherapy, etc. Typically, SYLARAS is deployed in three stages. In the first stage, longitudinal immunophenotyping data are collected from multiple mouse lymphoid organs of test and control subjects using an approach such as multiplex flow cytometry (FC). In the second stage, raw FCS files are spectrally compensated, filtered for viable cells and then stratified into distinct immune cell classes via graphical user interface (GUI)-assisted manual gating or clustering via PhenoGraph^17^ or FlowSOM^18^. In a final stage, data-rich graphical dashboards are generated, one per immune cell type as means of summarizing basal immune status and response to perturbation.

We demonstrate the use of SYLARAS by studying the impact of intracranially engrafted GL261 mouse glioma on peripheral immune cell composition in five major lymphoid organs. Although highly multiplexed methods for immune system profiling such as mass cytometry (CyTOF) are capable of measuring large numbers of features per cell (tens to hundreds), the method is relatively expensive. FC retains an advantage when large numbers of samples must be analyzed due to the speed, robustness, wide availability, and low cost of multiplex FC. The resource described in the current paper comprises, (i) a well-validated 12-channel FC panel able to distinguish major murine immune cell types, (ii) immuno-profiles for ~1×10^8^ cells from 240-tissues in control and GBM-bearing mice, (iii) Python-based SYLARAS software for the programmatic identification of tumor-induced changes in systemic immune composition.

As one illustration of the utility of this data, we study a previously undescribed change in immune architecture caused by GBM that involves a specialized subset of CD8^+^ T lymphocytes characterized by expression of the CD45R/B220 isoform of the CD45 protein tyrosine phosphatase type C receptor (*Ptprc*; henceforth B220^+^ CD8T cells). The previously described ability of B220^+^ CD8T cells to attenuate immune response to self-antigens^19,20,21,22^ suggests they may have immunosuppressive activity in GBM. We find that these cells are depleted from the circulation of tumor-bearing animals and infiltrate the TME of both mice and human gliomas. Relative to conventional CD8^+^ T cells, murine B220^+^ CD8T cells express different genes, are morphologically distinct, and localize to different regions of the brain tumor mass, suggesting that they represent a functionally distinct lymphocyte population. Experimental protocols, datasets, and source code for the SYLARAS project are freely-available via GitHub (https://github.com/gjbaker/sylaras), Synapse (https://www.synapse.org/#!Synapse:syn21038618), and links at the SYLARAS project website (https://www.sylaras.org).

## RESULTS

### A visual compendium of the peripheral immune response to GBM

To collect a dataset on the effects of GBM on the peripheral immune system, we harvested blood, bone marrow, deep/superficial cervical lymph nodes, spleen, and thymus from 48 age-matched, immunocompetent C57BL6/J mice engrafted with syngeneic GL261 glioma cells (National Cancer Institute Tumor Repository) or vehicle alone at three time points post-engraftment (7-, 14-, and 30-days; n=8 mice per treatment per time point). Brains were also collected at the time of euthanasia for t-CyCIF^23,24^: a method in which multiplex images are assembled from iterative rounds of conventional four-color immunofluorescence. Stereotactic injection of 3×10^4^ GL261 cells into striatum generated rapidly growing tumors with a median survival of 36 days (**Supplementary Fig. 1**). Immune tissues were disaggregated and immunolabeled with an optimized panel of 11 fluorophore-conjugated antibodies and then analyzed by FC (**Fig. 1** and **Supplementary Figs. 2 and 3**). Average cell viability was 98.6% (range: 90% - 99.7%) across the ~10^8^ cell dataset (**Supplementary Fig. 4a**). Immune cell abundance varied across the five lymphoid organs. To correct for these differences, a weighted random sample (WRS) of 1×10^7^ cells was drawn from the full dataset based on the cellularity of each tissue. This normalized the number of cells per tissue sample to an average of ~ 4×10^4^ (**Supplementary Fig. 4b**).

**Figure 1.**
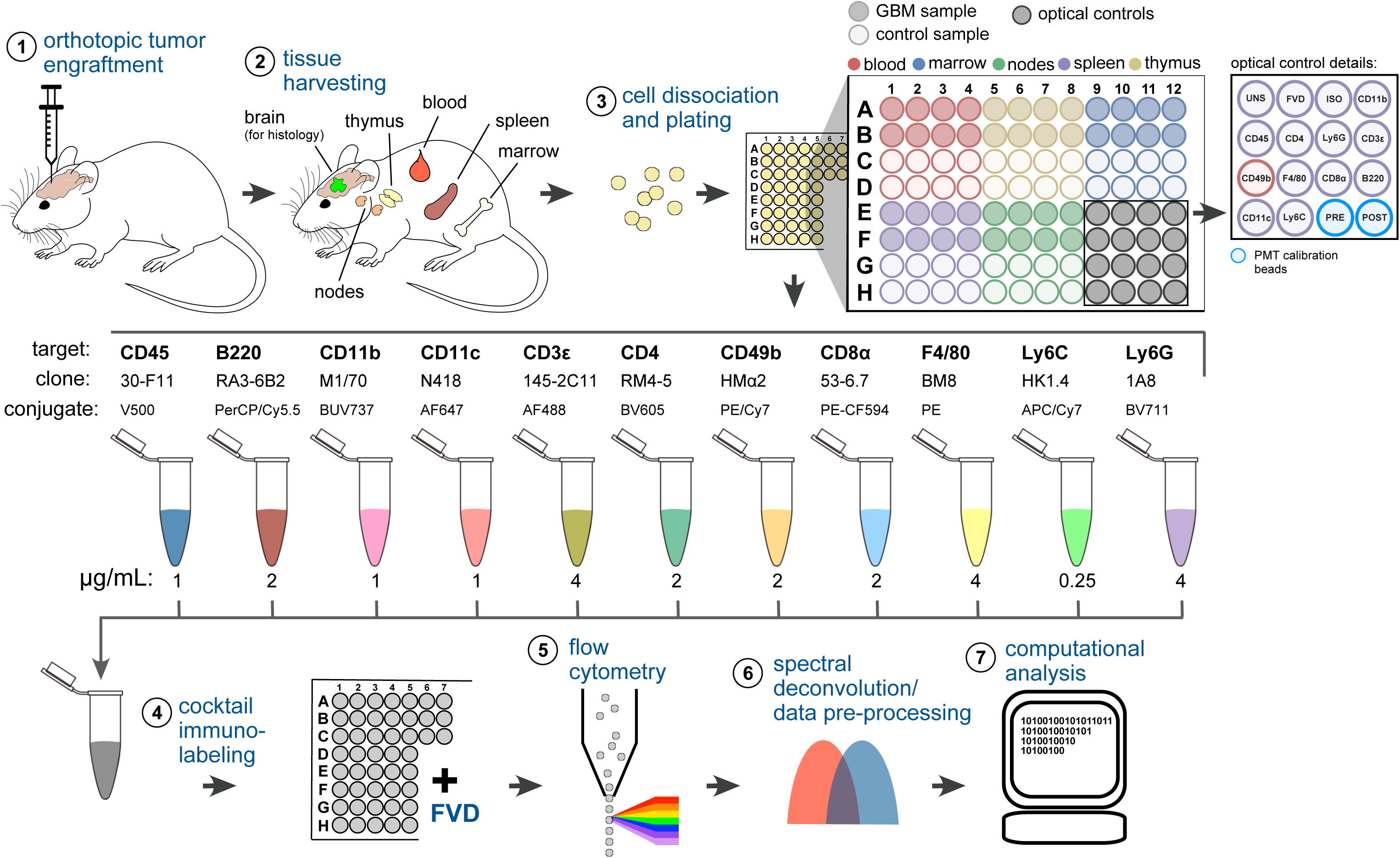
Immunoprofiling GBM-bearing mice by 12-color FC. (**1**) GBM cells (or vehicle control) were stereotactically engrafted into the striata of age-matched mice. (**2**) Lymphoid tissues were harvested from 8 replicate mice of each group 7-, 14-, and 30-days after tumor implantation. (**3**) Tissues were disaggregated and plated in a 96 well V-bottom plate. Locations of optical controls collected during each data acquisition are indicated. UNS=unstained splenocytes, FVD=fixable viability dye only, ISO=CD45 isotype control plus FVD, PRE=PMT calibration beads run before data acquisition, POST=PMT calibration beads run after data acquisition. (**4**) Cells were immunolabeled with 11 fluorophore-conjugated antibodies then stained with FVD. (**5**) Single-cell data were acquired by high-throughput FC. (**6**) Raw data were spectrally deconvolved and selected for viable singlets via conventional methods. (**7**) Preprocessed data underwent a histogram gating procedure (as described in Fig. 3) prior to computational analysis with SYLARAS.

To provide an overview of the 240-tissue dataset, we used SYLARAS to generate a set of graphical dashboards, each of which captured the statistics of a specific immune cell subset (**Fig. 2** and **Supplementary Fig. 5**). Dashboards specified the cell type alias (e.g. PMN for polymorphonuclear neutrophils), lineage (myeloid vs. lymphoid), immunomarker signature (e.g. CD45^+^ CD11b^+^, LyC^+^, Ly6G^+^), distribution among 5 lymphoid organs, and fractional contribution to all cells in the dataset (**Fig. 2a-e**). Quantitative information was also provided on light scattering and immunomarker profiles (**Fig. 2f,g**), mean differences and fold-changes between age-matched control and tumor-bearing mice across three time points in tumor progression (**Fig. 2h,i**), as well as cell type frequency in each tissue for every animal in the study (**Fig. 2j**). The modularity of the SYLARAS algorithm allows dashboard features to be changed by minor rewriting of the source code.

**Figure 2.**
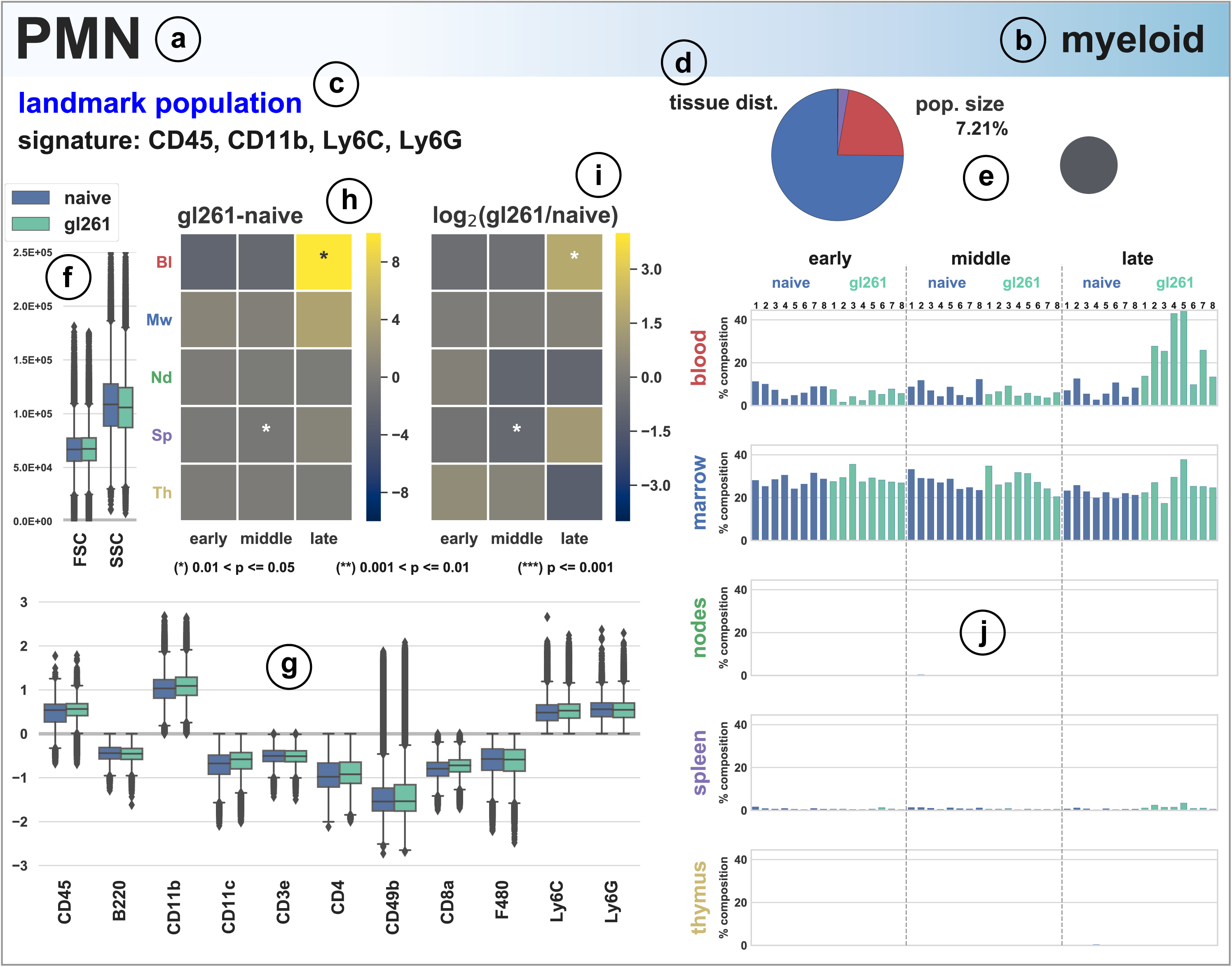
SYLARAS dashboard summarizing a GBM-induced immune system perturbation. Example SYLARAS dashboard for polymorphonuclear (PMN) immune cells displaying 9 cell type-specific attributes: (**a**) alias; (**b**) lineage; (**c**) immunomarker signature, “landmark population” indicating whether to immunophenotype corresponds to 1 of 14 major immune cell classifications; (**d**) distribution of cells across 5 lymphoid tissues color-coded as in (**h, i** and **j**); (**e**) percentage of all cells (in the random sample subjected to detailed analysis); (**f**) FSC/SSC; (**g**) Logicle-transformed, background-subtracted immunomarker signal intensity; (**h** and **i**) time and tissue-specific difference in mean percentage and log_2_ fold-change between GBM-burdened and mock-engrafted animals (n=8 mice/group), asterisks denote one of three levels of statistical significance; (**j**) percentage of each tissue across the study’s 48 mice.

### Manual and automated approaches to cell subset identification with SYLARAS

Manual gating of cytometry data is based on prior-knowledge about patterns of CD (cluster of differentiation) antigen expression and is usually performed manually using software programs such as FlowJo. However, the 2,640 fluorescence intensity vs. cell count histograms in our study (48 mice × 11 immunomarkers × 5 tissues) required the development of a more efficient approach. We therefore represented each histogram on a Logicle scale^25^ (a generalization of hyperbolic sine functions widely used to display compensated flow-cytometry data) and formatted as a scrolling HTML table viewed in a Web browser. This facilitated comparison between tissues, time points, and replicates, enabling rapid identification and recording of gate values between positive and negative signal intensities (**Fig. 3a**). A spreadsheet of these values was fed into SYLARAS. Gate values were then subtracted from intensity distributions to center the gate at zero and make non-specific fluorescence negative valued (**Fig. 3b**). Gating the data in this way, we were able to process the entire dataset in <2 hours. Peak finding algorithms and Gaussian mixture models^26^ were evaluated as a means to automatically setting gates, but provided no significant advantage over manual curation when the time for human review was included. Following the gating procedure, SYLARAS was used to binarize intensities according to mathematical sign, resulting in an M-dimensional Boolean immunophenotype for each cell where M is the number of immunomarkers used in the analysis (e.g. CD_1_^+^, CD_2_^−^, CD_3_^+^… CD_M_^−^) (**Fig. 3c**).

**Figure 3.**
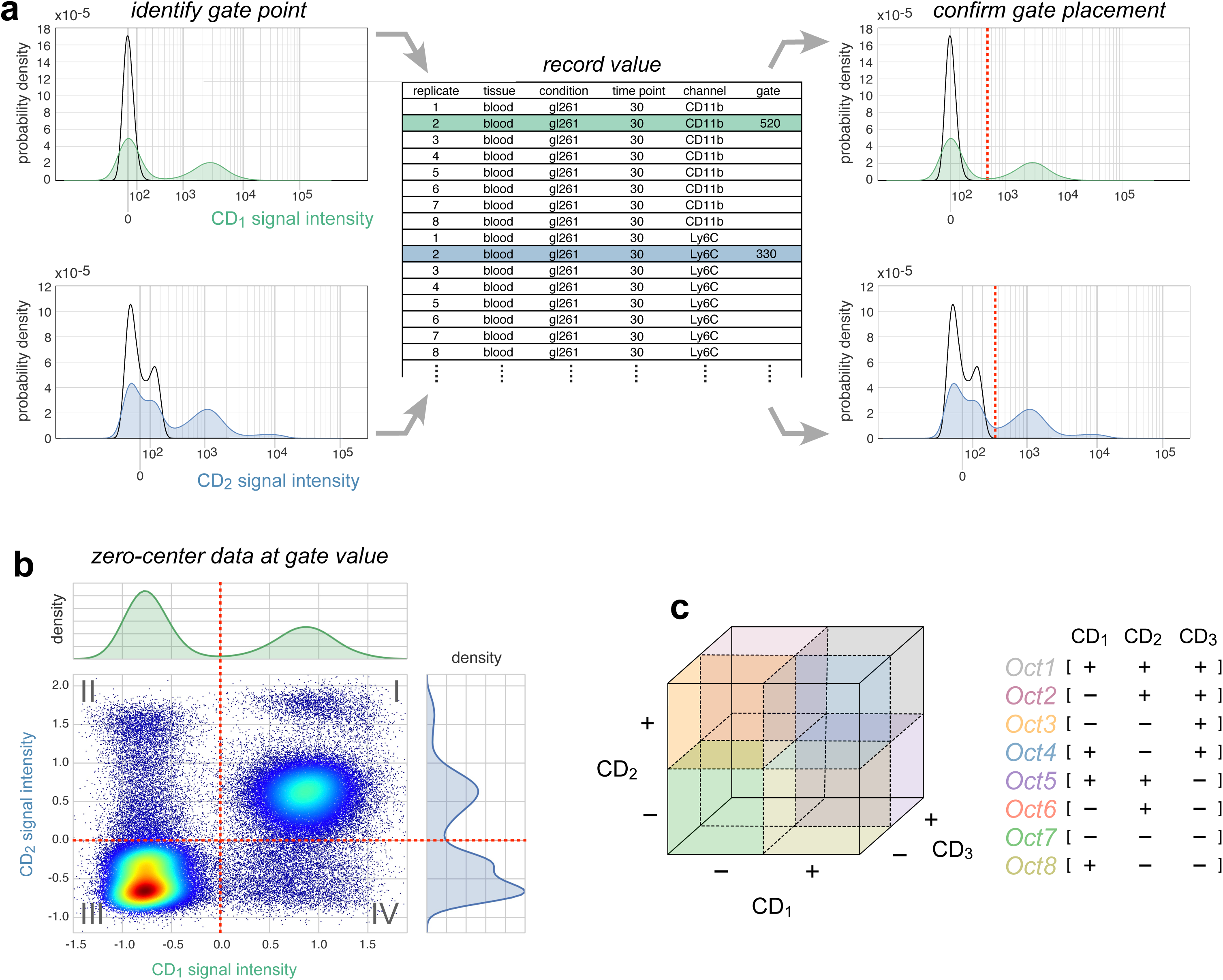
The SYLARAS approach to immune cell subset identification. (**a**) Illustration of an iterative procedure in which intensity vs. cell number histograms were gated to distinguish positive staining signal and autofluorescence. Two example histograms are shown (CD_1,_ green; CD_2_, blue). Black outlines superimposed on each histogram represent the signal intensity distribution of unlabeled (control) splenocytes which SYLARAS includes as a fiducial reference for background signal intensities. Gate values are entered into a .TXT (or .XLS) file preformatted with sample metadata which, once complete, is supplied as input into a program that updates the histograms with a dashed red line at the curated gate for confirmation or refinement. (**b**) Bivariate scatter plot showing the same signal intensity data as shown in panel (a) after Logicle-transformation and subtraction of respective gate values from the signal intensity of every cell in the distribution demonstrating the binning of cells among 2^2^=4 possible immunophenotype quadrants. (**c**) Extension of quadrant binning to 3-dimensional data where cells are binned among 2^3^=8 possible immunophenotype octants.

Of the 2^11^ (2,048) Boolean immunophenotypes that can be specified with 11 markers, 604 were represented by ≥1 cell, and 30 were populated by >1% of cells in one or more of the 240 tissues analyzed. Together these 30 immunophenotypes accounted for 97% of viable immune cells and were the focus of further analysis. The 30 immunophenotypes were divided into 14 landmark immune cell subsets based on marker expression (e.g. CD4^+^ or CD8^+^ T cells, etc.; **Fig. 4a**, inner wedges). Landmark populations were further divided into subclasses based on expression of additional immunomarkers. For example, CD8^+^ T cells characterized as CD45^+^, CD3ε^+^, and CD8α^+^, which also expressed Ly6C, correspond to Ly6C^+^ CD8^+^ T cells, which are mouse memory T cells^27^ (**Fig. 4a**, outer wedges). Relative to all cells scored, the abundance of cells in the 30 immunophneotypes ranged from ~30% for B cells to ~0.01% for dendritic cells (DCs) (**Fig. 4b**). In agreement with the established light scattering properties of different immune cell subsets, forward and side scatter (FSC and SSC) were lowest among cells classified as adaptive lymphocytes (e.g. CD4^+^ and CD8^+^ T cells), intermediate in non-polymorphonuclear myeloid cells and innate lymphocytes (e.g. monocytes/macrophages, DCs, and NK cells), and highest among granulocytes (e.g. neutrophils) (**Fig. 4c**). The exceptionally high SSC of a subset of CD45^+^, CD11b^+^, F4/80^+^ cells allowed them to be classified as eosinophils (as opposed to macrophages).

**Figure 4.**
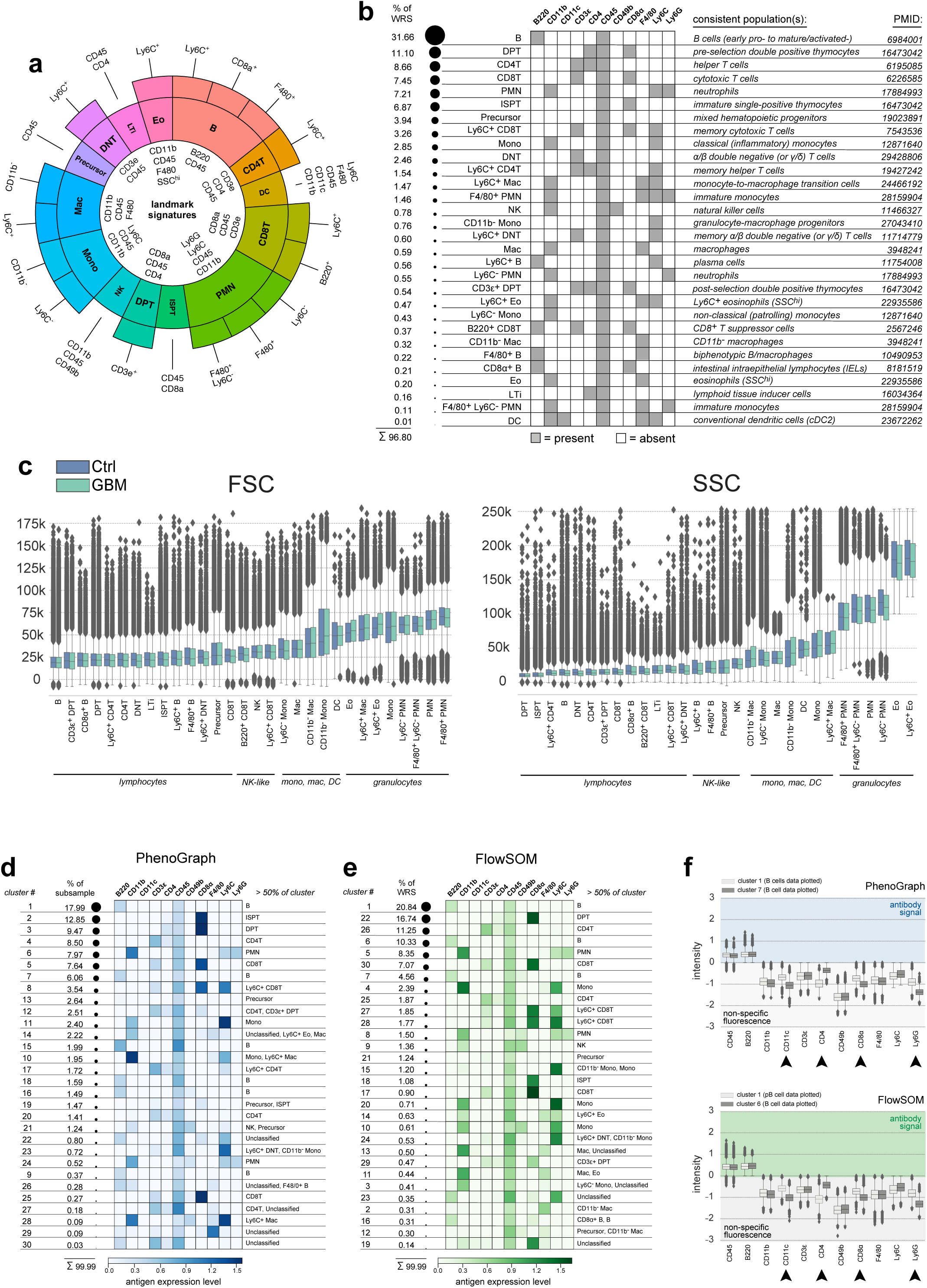
Mouse Immune cell subset identification via knowledge-based and data-driven methods. (**a**) Mapping of immunophenotypes to known cell lineages. Fourteen landmark immune cell populations are indicated by the inner wedges of the sunburst plot. Outer wedges represent refinements within the landmark populations. Immunomarkers expressed by each cell type are specified. (**b**) Fraction of the WRS accounted for by each cell type (left), their immunophenotype (middle), and the known immune cell subset(s) consistent with each (right, PMID=PubMed identifier). (**c**) FSC and SSC distributions of cells from control (blue boxplots) and GBM-burdened mice (green boxplots) annotated to each cell type and shown in ascending rank order from left-to-right according to the median value of the control data. Box plot elements: horizontal line, median; box limits, first to third quartile (Q1 to Q3); whiskers, from Q1-1.5 × interquartile range (IQR) to Q3 + 1.5 × IQR; diamond points, outliers. (**d** and **e**) The fraction of data accounted for by each of 30 PhenoGraph (*k*=20, Euclidean distance, left panel) and 30 FlowSOM (*nClus*=30, right panel) clusters. Heatmaps show mean immunomarker signal intensity of cells constituting each cluster. Background signal intensities were set to zero to highlight foreground antibody signal. Cell type calls made during knowledge-based cell subset identification accounting for >50 of cells in each cluster are indicated to the right of each heatmap. (**f**) Logicle-transformed signal intensity distributions of PhenoGraph (top) and FlowSOM (bottom) cluster pairs enriched for CD8T cells demonstrating population splitting based on on-target signal intensities. Signal intensities above zero indicate foreground antibody signal while negative values indicate non-specific fluorescence as determined by our histogram gating procedure. Arrowheads indicate immunomarker channels exhibiting considerable difference between clusters. (**g**) Logicle-transformed signal intensity distributions of PhenoGraph (top) and FlowSOM (bottom) cluster pairs enriched for B cells demonstrating population splitting based on non-specific signal intensities.

Unsupervised clustering is a common way to identify immune cell types in large datasets since it circumvents the need for gating^28^. We integrated two widely used clustering algorithms into the SYLARAS pipeline: PhenoGraph^17^, which is based on nearest-neighbor clustering, and FlowSOM^18^, which uses a minimum spanning tree algorithm. Because of the computational burden of PhenoGraph, random sampling was used to reduce the dataset a further 5-fold. To facilitate comparison between clustering methods and immune cell classification by gating, the number of nearest neighbors (*k*) in PhenoGraph and number of metaclusters (*nClus*) in FlowSOM were adjusted so that both algorithms generated 30 clusters: the same number of cell subsets identified manually. Unexpectedly, no clusters generated by either method were comprised entirely of cells identified by manual gating as a single immunophenotype. The best agreement was obtained for 19 PhenoGraph clusters and 21 FlowSOM clusters in which >50% of clustering cells were assigned a single immunophenotype (**Fig. 4d,e** and **Supplementary Fig. 6**). For an additional 8 PhenoGraph and 7 FlowSOM clusters, cell populations were made up of two to three immunophenotypes, while the remaining clusters had no obvious correspondence with a known immune cell subset.

Why are results obtained by clustering algorithms so different from those from manual gating? In some cases, it was clear that cells of the same manually assigned immunophenotype were split into multiple clusters based solely on differences in intensities values corresponding to background autofluorescence as determined by human inspection. PhenoGraph clusters 1 and 7 and FlowSOM clusters 1 and 6 were good examples of this phenomenon; cells in these clusters expressed similar levels of CD45 and B220 (an immunomarker of B cells) but differed in background florescence in channels used for AF647, BV605, PE-CF594, BV711 (corresponding to CD11c, CD4, CD8α, and Ly6G respectively) (**Fig. 4g** and **Supplementary Fig. 6**). These background intensities were at least 10-100-fold lower than CD11c, CD4, CD8α, and Ly6G staining on other cells in the dataset (e.g. CD4 on T cells). By manual gating, low signal intensities are simply assigned a value of zero; in contrast, automated clustering identifies small absolute differences in background as statistically significant, and uses them to assign cluster membership. We speculate that, with a large number of relatively low dimensional FC histograms, manual gating—which effectively represents a fully supervised method— may generally be superior to unsupervised clustering.

### Impact of GBM on peripheral immune cell frequency and network-level architecture

The composition of five lymphoid organs with respect to the 30 immunophenotypes identified by manual gating was as expected: the spleen, cervical lymph nodes, and blood were primarily made up of B cells, CD4^+^ T cells, and CD8^+^ T cells, the bone marrow was primarily composed of PMN cells, B cells, and monocytes/macrophages, and the thymus was predominantly double-positive T (DPT) cells and immature single-positive T (ISPT) cells (**Fig. 5a,b**). When the data were analyzed by principle component analysis (PCA) we found that the first two principle components (PCs) explained >60% of dataset variation—a good performance for PCA. The scores plot for these 2 PCs revealed the presence of five primary clusters separated by tissue type (**Fig. 5c**). As expected, spleen, lymph nodes, and blood exhibited substantial overlap since the cellular composition of these secondary lymphoid tissue is more similar then that of primary lymphoid organs (i.e. bone marrow and thymus). GBM-burdened mouse 3 at t=30-days was an outlier (**Fig. 5c**). This mouse had more DCs, macrophages, and Ly6C^−^ PMN cells in its blood and bone marrow relative to any other GBM-burdened animal at the 30-day time point (**Fig. 5d**).

**Figure 5.**
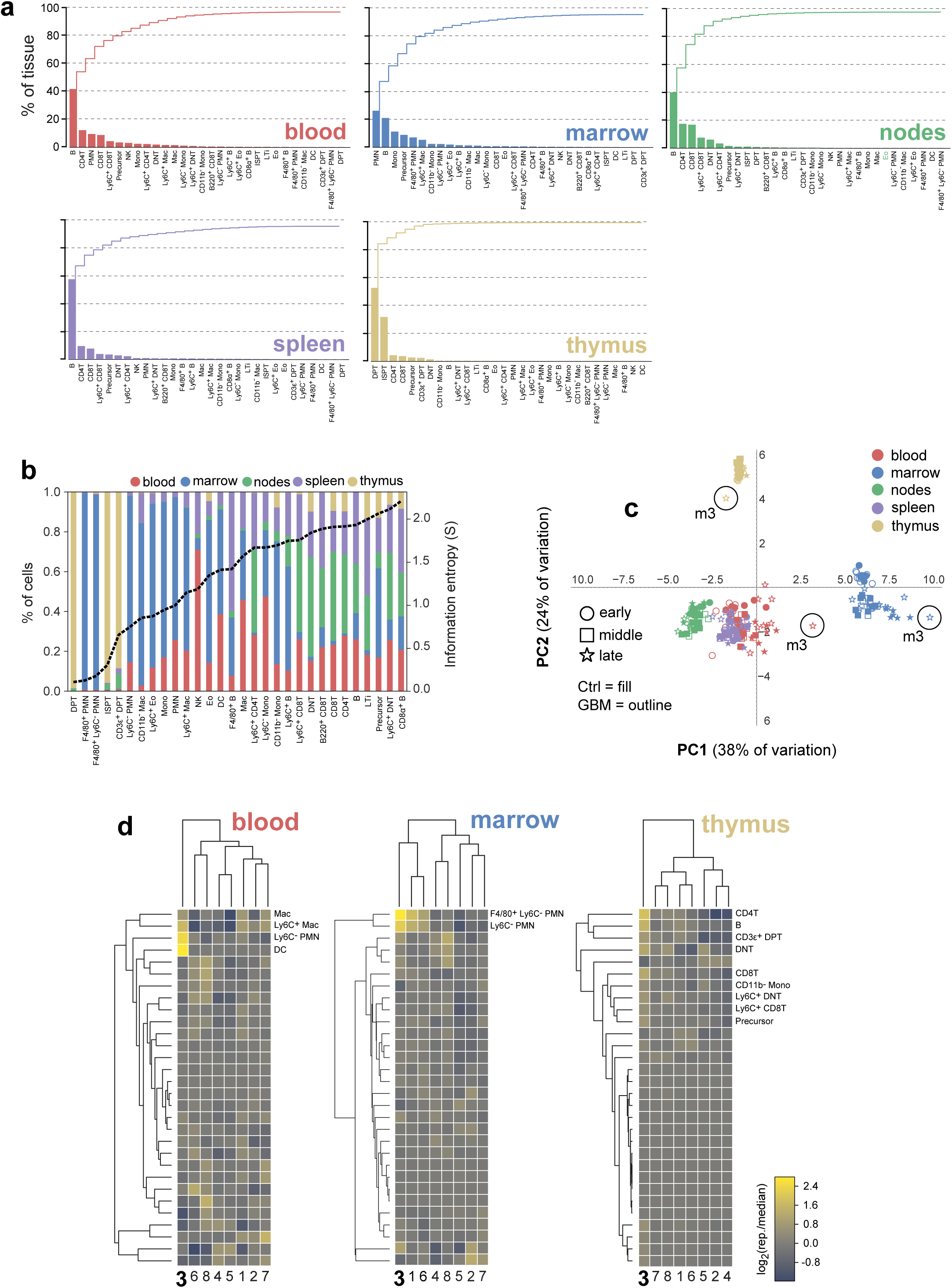
Breakdown of mouse lymphoid tissue composition in 30 cellular immunophenotypes. (**a**) Plots of the individual (bars) and cumulative (stairs) percentage of 5 lymphoid tissues accounted for by successively scarce cell types. (**b**) Distribution of cell types among lymphoid tissues (left y-axis) shown in ascending rank order from left-to-right according to their information entropy (dashed black line, right y-axis). DPT and ISPT cells had the lowest entropy due to their near-exclusive restriction to the thymus, while mature CD4^+^ T and CD8^+^ T cells had the highest entropy due to their broad distribution across multiple tissues. (**c**) Scores plot of the first 2 PCs of a PCA performed on a 240-row × 30-column data table of the percentage of each tissue sample (rows) accounted for by each cell type (columns). Circles highlight samples from GBM-burdened mouse 3 at t=30-days. (**d**) Clustermaps of the log_2_ ratio of the tissue-specific frequency of each cell type in late-stage GBM-burdened mice relative to the median value for the group of 8 mice. Cell types contributing most strongly to mouse 3’s outlier tendency are indicted per tissue. Clustering was performed using Euclidean distance and the unweighted pair group method with arithmetic mean (UPGMA) linkage method.

To identify time and tissue-dependent differences in immunophenotypes between healthy and disease-burdened animals we performed 450 two-tailed Student’s t-tests (30 immunophenotypes × 5 tissues × 3 time points). After adjusting for multiple hypothesis testing using the false discovery rate, 25 statistically significant differences were identified (q-value <0.05; **Fig. 6a**), most of which were observable at t=30-days. However, increases in the abundance of double-negative T (DNT) cells in the spleen and lymph nodes of tumor-bearing mice relative to controls were observed as early 7-days post-engraftment. This likely reflects the immune system’s initial response to tumor cell inoculation as opposed to frank tumor burden and is consistent with the role played by DNT cells in the acute response to inflammation^29^.

**Figure 6.**
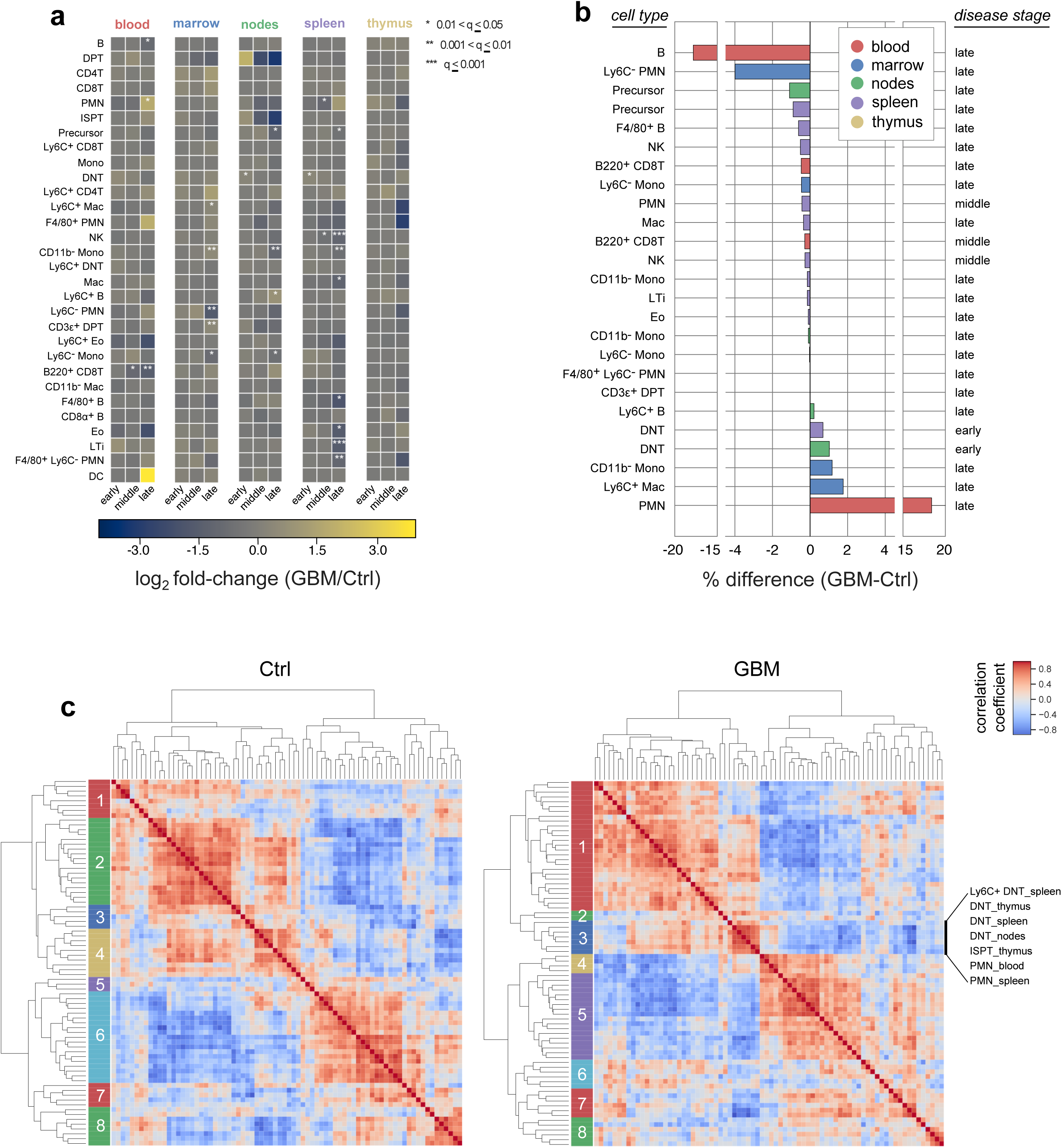
GBM perturbs peripheral immune cell subset frequency and correlation structure. (**a**) Average log_2_ fold-change in time and tissue-specific cell type frequency between GBM-bearing and mock-engrafted mice. Asterisks denote one of three levels of statistical significance. (**b**) Average percent change in time and tissue-specific cell type frequency between GBM-bearing and mock-engrafted mice. Only statistically significant cell subsets are shown (q≤0.05). (**c**) Agglomerative hierarchical clustering of Spearman’s rank-order correlation coefficients computed across replicates of control (left) and GBM-burdened (right) mice. Clustering was performed using the UPGMA linkage algorithm and Euclidean distance. Members of GBM-burdened cluster 3 are specified in order from top-to-bottom.

Tumor-associated decreases in immune cell frequency were observed for circulating B cells, splenic lymphoid tissue inducer (LTi) cells^30^, splenic biphenotypic B/macrophages^31^, CD45^+^ Lin^−^ immune progenitor cells in the lymph nodes and spleen, and Ly6C^−^ PMN cells in the bone marrow. Tumor-associated increases were also observed. CD3ε^+^ post-selection thymocytes accumulated in the bone marrow of tumor-bearing mice. This result was consistent with the work of Chongsathidkiet et al. showing that GBM-induced T cell lymphopenia is in part due to sequestration of cells in the bone marrow^32^. Ly6C^+^ plasma cells^33^ were also increased in the cervical lymph nodes, suggestive of an active humoral immune response and consistent with the known ability of brain-derived antigens to traffic to the cervical lymph nodes and elicit plasma cell responses^34^. Myeloid cells also changed in response to tumor-burden: for example, increases in circulating CD11b^+^ Ly6C^+^ Ly6G^+^ PMN cells, consistent with granulocytic myeloid derived suppressor cells (MDSCs), were observed. These cells are known to accumulate in the blood of GBM-burdened mice and humans^35,36^ and were 18% more abundant on average in diseased animals at t=30-days (**Fig. 6b**). Ly6C^−^ patrolling monocytes, eosinophils, and many other myeloid populations also increased in some tissues while decreasing in others. All of these changes are detailed in supplementary materials and summarized in the data available through SYLARAS dashboards (**Supplementary Fig. 5**).

Variation in the frequencies of immune cells in different tissues among mice of the same treatment group suggested a way to investigate immune system homeostasis. Between-subject biological variation (BSBV)^37^ is a potential confounder in some statistical tests due to increasing data variance and correspondingly weaker p-values; however, correlated variation, as determined by correlation analysis, may suggest functional interaction between cell types. The effect of GBM on correlated variation between peripheral immune cells was investigated by computing Spearman’s rank-order correlation independently on the tissue-specific immunophenotypes of healthy and diseased animals. Cell populations with a frequency <0.1% were excluded from the analysis to minimize noise leaving 76 tissue-specific immunophenotypes for which correlations were performed. Agglomerative hierarchical clustering on the resulting correlation matrices revealed 8 clusters per treatment group as determined by silhouette analysis (**Fig. 6c**).

Cell-cell correlations for GBM-burdened and control mice were substantially different. For example, GBM-burdened mice exhibited a cluster (cluster 3) significantly enriched for DNT cells in the spleen, thymus, and draining lymph nodes (p=0.0003) and PMN cells in the blood and spleen (p=0.04) that was not present in control data, suggesting that GBM induces coordinated changes in these two immune cell subsets across multiple tissues. The results of Coffelt et al.^38^ support this finding; they show that IL-17A-producing γδ T cells—which likely correspond to DNT cells in our study—induce a systemic inflammatory cascade leading to the accumulation of circulating PMN neutrophils in a mouse model of breast cancer. More generally, we propose that correlation analysis can be used to probe the self-regulatory architecture of the peripheral immune system and its perturbation by disease. Our data show that peripheral immune architecture is broadly altered by GBM.

### B220^+^ CD8T cells are depleted in the blood of GBM-bearing mice and transcriptionally-distinguishable from other CD8^+^ T lymphocytes

Among the statistically significant differences in immune cell frequency that were detected between healthy and tumor-bearing animals, two cell populations stood out as exhibiting monotonic changes with increasing statistical significance across time: splenic NK cells and circulating B220^+^ CD8T cells. Both populations decreased from t=14-days (q=0.017 for NK cells; q=0.026 for B220^+^ CD8T cells:) to t=30-days (q=0.0009 for NK cells; q=0.002 for B220^+^ CD8T cells) relative to age-matched control mice. NK cells have previously been associated with GBM^7,8,39^ but the disease has not previously been shown to involve B220^+^ CD8^+^ T lymphocytes, cells known to attenuate immune responses to self-antigens in mice and humans.^19,20,21^

B220, also known as CD45R, is an isoform of the *Ptprc* gene expressed by murine B cells at all developmental stages from pro-B cells through mature B cells. Unlike the uniformly low B220 staining intensity in CD4^+^ T cells and high staining in B cells—the canonical B220-expressing immune cell type—B220 intensity in CD8^+^ T cells spanned several orders of magnitude; ~5-10% of CD8^+^ T cells expressed B220 at levels similar to those of B cells (**Fig. 7a**). Three lines of evidence suggested that these B220^hi^ CD8^+^ T cells represent a distinct cell subset from conventional CD8^+^ T cells. First, whereas the abundance of circulating B220^+^ CD8T cells decreased with advancing disease, CD8^+^ T cells were unaffected (**Fig. 7b**). Second, B220^+^ CD8T cells were remarkably rare in the thymus, but more abundant in the bone marrow and spleen given their fractional contribution to total CD8^+^ T cell counts (**Fig. 7c**). Third, although the frequency of splenic B220^+^ CD8T and CD8^+^ T cell was strongly correlated in most mice (R^2^ between 0.6 and 0.9), this correlation was lost in late-stage, disease-bearing animals (R^2^=0.01), suggesting a differential response to the presence of GBM (**Fig. 7d**).

**Figure 7.**
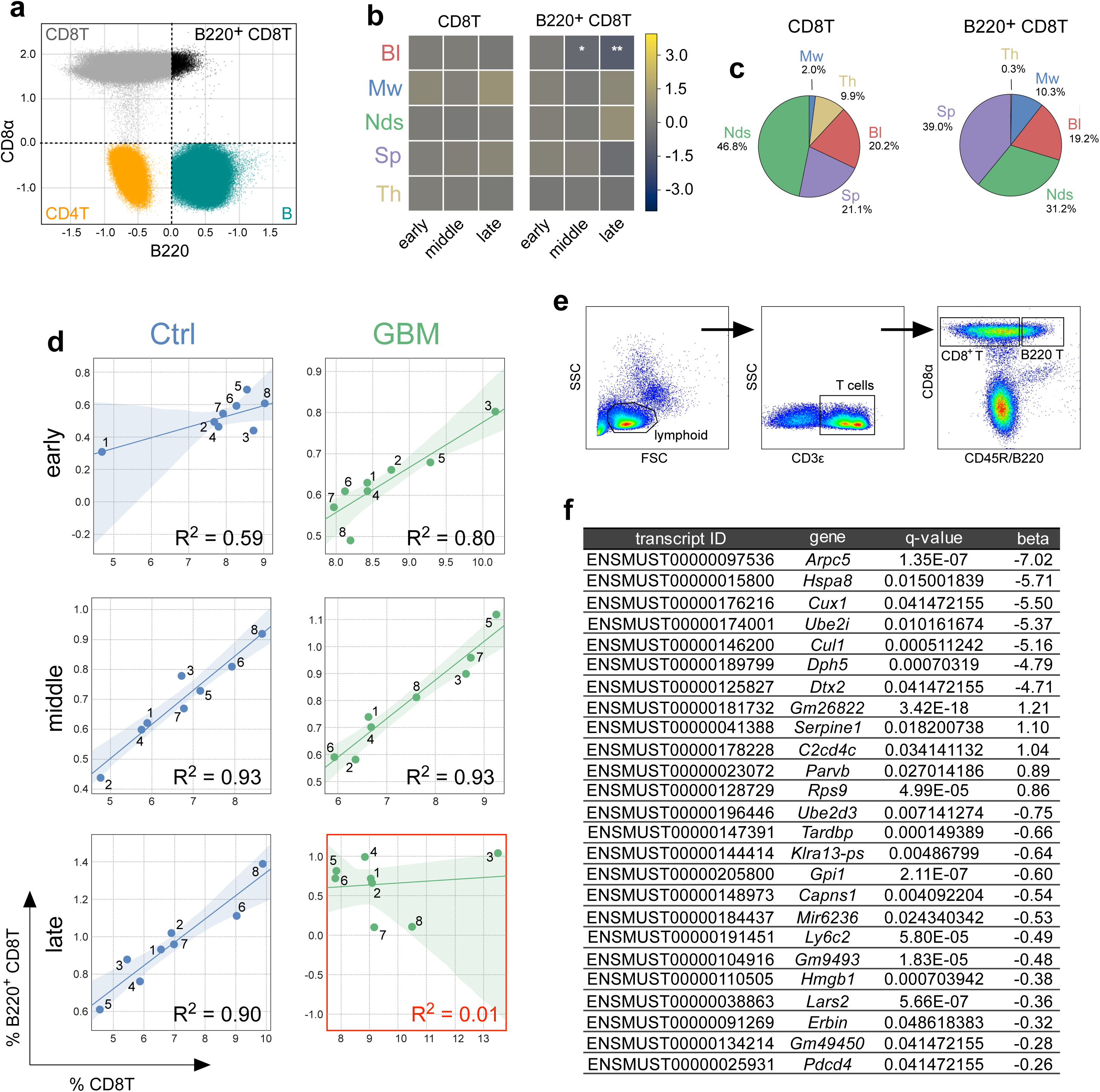
B220^+^ CD8T cells are transcriptionally-distinct from other CD8^+^ T lymphocytes. (**a**) Logicle-transformed B220 (x-axis) vs. CD8α (y-axis) expression of cells classified (clockwise from upper-left quadrant): CD8^+^ T, B220^+^ CD8^+^ T, B, and CD4^+^ T cells. (**b**) Average log_2_ fold-change in time and tissue-specific cell type frequency between GBM-bearing and mock-engrafted mice for the CD8T (left) and B220^+^ CD8T (right) cell types. (**c**) Distribution of CD8T (left) and B220^+^ CD8T (right) cells across the 5 lymphoid tissues analyzed in this study. (**d**) Regression plots showing the frequency of splenic CD8T cells (x-axes) versus splenic B220^+^ CD8T cells (y-axes) from control (left column) and GBM (right column) mice across the study’s 3 time points (rows). Data points represent 1 of 8 replicate mice. Regression plot elements: line, regression line; transparency, 95% confidence interval for the regression; R^2^, coefficient of determination. (**e**) Gating strategy for the FACS-purification of B220^hi^ and B220^lo^ CD8^+^ T cells from the CD19-immunodepleted spleens of normal C57BL/6J mice. (**f**) Table of differentially expressed transcripts between B220^hi^ and B220^lo^ CD8^+^ T cells sorted by the absolute value of their beta value (effect size on log-transformed data).

To show directly that B220^+^ CD8T cells differ from other CD8^+^ T lymphocytes, we FACS-sorted total CD8^+^ T cells from the spleen of healthy mice into B220^lo^ (n=3) and B220^hi^ (n=2) fractions and performed RNA sequencing (**Fig. 7e**). Using the Sleuth RNA-seq data analysis tool^40^, we identified 25 transcripts that were differentially expressed between the two cell populations (Wald test, q-value ≤0.05) (**Fig. 7f**). The largest difference involved a splice variant encoding subunit 5 of the ARP2/3 complex (*Arpc5*, transcript id: ENSMUST00000097536), which was 7-fold lower in B220^hi^ cells compared to their B220^hi^ counterparts. This gene promotes actin polymerization^41^ and maintenance of T cell receptors at the plasma membrane^42^. Other differentially expressed genes have been associated with protein synthesis (*Cul1*, *Dph5*, *Lars2*, and *Rps9*) and proteasome-mediated protein degradation (*Ube2d3*, *Dtx2*). We conclude that B220 expression by CD8^+^ T cells is associated with an altered transcriptional state, differential tissue distribution under normal circumstances, and selective response to the presence of GBM.

### B220^+^ CD8T cells infiltrate mouse and human brain tumors

We postulated that the observed reduction in circulating B220^+^ CD8T cells in tumor-bearing mice might be a consequence of their extravasation from the circulation into the brain TME. We therefore characterized tumor-infiltrating immune cells using multiplex t-CyCIF. An antibody panel functionally similar to the one used for FC was optimized and validated (**Supplementary Fig. 7** and **Supplementary Table 1**) and then used to acquire 12-channel images of the late-stage brain TME. One-hundred sixty-eight (168) fields of view (tiles) were collected at 40x magnification then registered and stitched together to form a composite image from which protein expression data could be extracted following image segmentation of ~9 x10^4^ singe cells comprising the tumor mass (**Fig. 8a** and **Supplementary Fig. 8a**). Using the immune cell identification features in SYLARAS, we identified multiple tumor-infiltrating immunophenotypes, including B220^+^ CD8T cells (**Fig. 8b** and **Supplementary Fig. 8b-d**). Relative to lymphocytes single-positive for CD8, the double-positive cells had a distinctive morphology: they were more eccentric, their plasma membranes were smoother, and their CD8α staining was more intense (**Fig. 8c** and **Supplementary Fig. 8b,c**). Their higher CD8α staining was consistent with data on peripheral B220^+^ CD8T cells, for which median CD8α staining was ~1.3-fold higher than other circulating CD8^+^ T cells (p<1×10^−4^; **Fig. 8d**). Tumor-infiltrating single-positive and double-positive lymphocytes also occupied different regions within the tumor mass: the former were more frequent at the tumor borders in areas of low tumor cell density, whereas double-positive cells were more evenly distributed in the center of the tumor (**Fig. 8e,f**).

**Figure 8.**
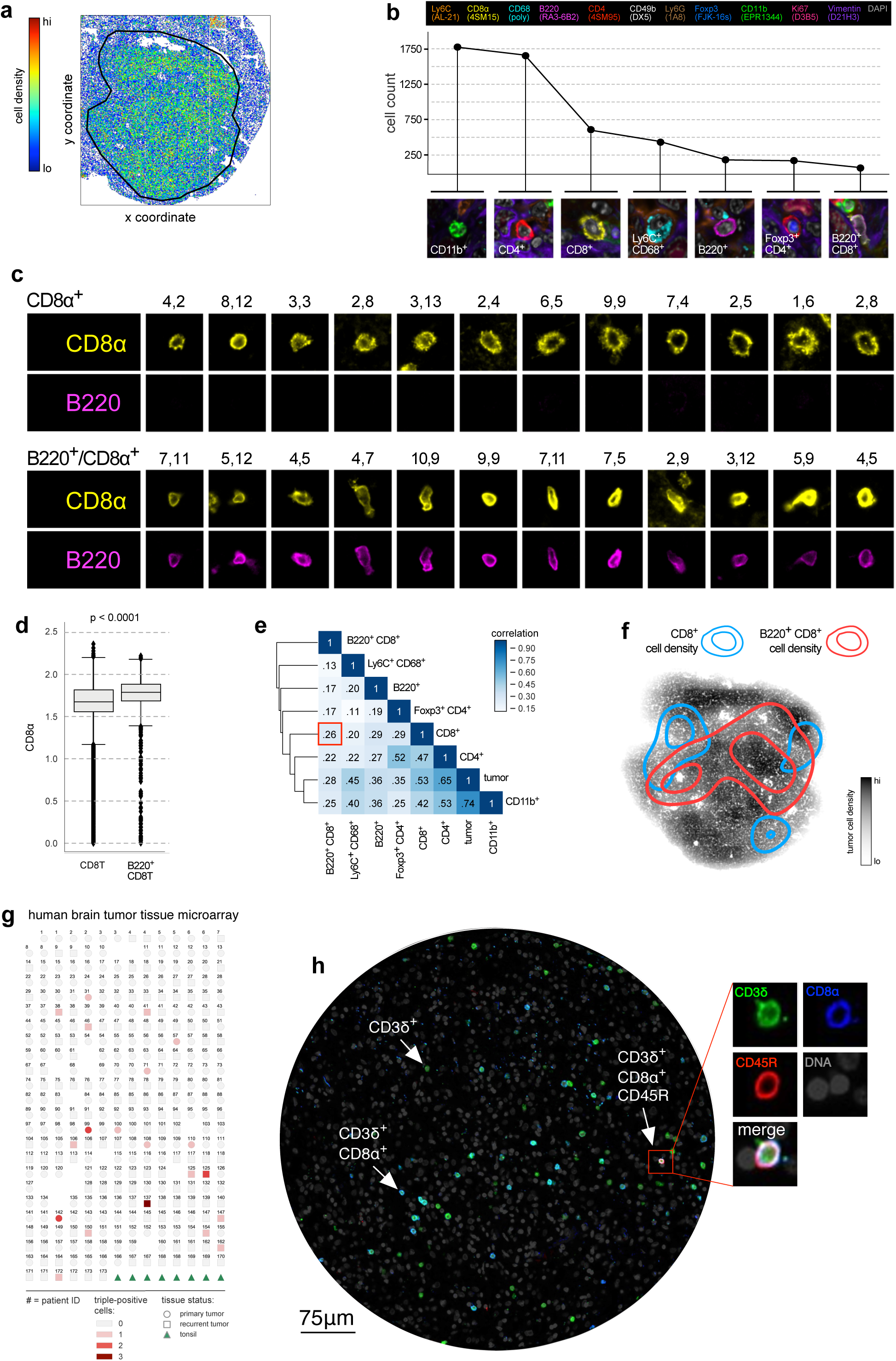
B220^+^ CD8T cells infiltrate the brain TME of mice and humans. (**a**) Spatial coordinates of ~9×10^4^ cells from the tumor-ipsilateral hemisphere of a GL261 GBM tumor 36-days after engraftment. Data points representing individual cells have been pseudo-colored according to density. Black perimeter outlines the tumor/brain parenchyma interface. (**b**) Cell counts and an example image of the 7 most abundant immunophenotypes in the brain TME. (**c**) Examples of CD8α single-positive (top) and B220/CD8α double-positive (bottom) lymphocytes identified throughout the brain TME. Tile coordinates of each cell are provided to allow for cross-referencing with panel (a) of Supplementary Fig. 8. (**d**) Box plot distributions of CD8α signal intensity for CD8T (left) and B220^+^ CD8T (right) cells in the blood of the study’s 48 mice. (**e**) Spatial correlation between immunophenotypes within the brain TME. Correlation between CD8α single-positive and B220/CD8α double-positive cells is highlighted by the red box. (**f**) Kernel density estimates of the spatial localization of CD8α single-positive (blue contour lines) and B220/CD8α double-positive (red contour lines) lymphocytes superimposed on a scatter plot of Vimentin^+^ Lin^−^ GBM cells shaded according to cell density. (**g**) Metadata map of a human brain tumor TMA containing 316 (0.6mm diameter) cores taken from gliomas resected from 173 different patients (numerical labels). The number of B220^+^ CD8T cell analogs (i.e. CD3δ^+^, CD8α^+^, CD45R^+^ cells) is indicated by the intensity of red shading. B220^+^ CD8T cells were identified in 19 cases, of which 16 cases were diagnosed as WHO grade IV primary or recurrent GBM, 3 diagnosed as grade III anaplastic oligodendroglioma or anaplastic astrocytoma and 4 were immunoreactive to antibodies recognizing the IDH1 (R132H) mutant (a key oncogenic driver in GBM). Ten (10) cases were male and nine female. Tumor status is as indicated: circles=primary tumor, squares=recurrent tumor, green triangles=human tonsil (positive control tissue for immunomarkers). (**h**) Core from an *IDH1* (R132H)-negative recurrent human GBM resection exhibiting a CD3δ^+^, CD8α^+^, CD45R^+^ cell (red square). CD3δ single-positive and CD3δ/CD8α double-positive T cells are also seen.

To determine whether cells analogous to mouse B220^+^ CD8T cells are present in human GBMs, we used human-specific antibodies targeting CD3δ, CD8α, and CD45R to immunolabel a tissue microarray (TMA) consisting of 312 glioma tissue cores resected from 173 different brain tumor patients (1-2 0.28 mm^2^ tissue cores per patient). The immunolabeled TMA was then scanned at 20x magnification to generate a 875-tile mosaic image (to view the full image, visit https://www.cycif.org/data/baker-2019/). Lymphocytes expressing all three immunomarkers (i.e. CD3δ^+^, CD8α^+^, CD45R^+^ T cells) were identified in 19 cores of different disease subtypes accounting for an average of 1.2% of all T cells detected (**Fig. 8g,h**). Thus, cells analogous to mouse B220^+^ CD8T cells are present in a variety of primary and recurrent clinical brain tumor specimens irrespective of sex or *IDH1* (R132H) mutational status.

## DISCUSSION

In this paper we describe a software tool for efficient analysis of the systemic immune system (SYLARAS) and a dataset collected using the tool that describes the impact of GBM on immune cells in multiple immune organs using a well-established syngeneic mouse model. Each of ~10^8^ cells from five primary and secondary immune organs was assayed for 11 CD antigens indicative of cell type and differentiation status as well as a viability and FSC and SSC. An 11-plex Boolean immunophenotype can specify 2,048 possible cell states, but we found that only 30 such states were populated by >1% of cells in any single tissue sample; across the entire data set, the frequency of these immunophenotypes ranged from 0.01% for CD45^+^, CD11b^+^, CD11c^+^, Ly6C, F4/80^+^ dendritic cells to 32% for CD45^+^, B220^+^ B cells. In aggregate, the cells described in our dataset comprised >97% of viable CD45^+^ cells. We observed substantial BSBV in these frequencies, even among age-matched and genetically-identical animals, and patterns of correlation and anti-correlation computed from this variability provides insight into the overall architecture of the immune system. Based on this, we find that cell frequencies in the spleen, thymus, blood, lymph nodes and bone marrow are broadly altered by GBM, demonstrating the disease imposes a broad, system-level perturbation of the immune system.

One GBM-dependent change in immune cell frequencies that we studied in greater detail involved an atypical subpopulation of CD8^+^ T cells expressing the CD45R/B220 splice variant of the phosphotyrosine phosphatase CD45. Across animals, conventional splenic CD8^+^ T and B220^+^ CD8^+^ T cells are highly correlated in abundance except in late-stage GBM-bearing animals. In these animals, B220^+^ CD8T cells are depleted from the circulation and accumulate in the GBM TME. These cells differ morphologically and transcriptionally from conventional CD8T cells and have been described previously as “CD8^+^ T suppressor cells”^21,22^. Although the functions of these cells are incompletely understood, activating them using monoclonal antibodies against the CD45R/B220 receptor increases proliferative responses to the mitogenic lectin PHA (phytohemagglutinin)^19,43^. B220^+^ CD8T cells are also capable of secreting the anti-inflammatory cytokines IL-10 and TGF-β^44^ and suppressing autoimmunity through direct cell-to-cell contact with self-reactive CD4^+^ T cells^21,22^. We find that human B220^+^ CD8T cells also accumulate in the TME of high-grade gliomas of patients, suggesting common properties for these cells in mice and humans. Whether the B220 antibodies used in our mouse studies (clone: RA3-6B2) recognize a functionally identical cell population as those used in our human studies (clone: MB1) remains uncertain; however, both antibodies are documented to react with restricted epitopes on exon A at the extracellular domain of a high molecular weight isoform of the CD45 glycoprotein expressed by B cells and a subset of T-cytotoxic and T-suppressor cells. Efforts are currently underway to establish the biological significance of B220^+^ CD8T cells in GBM and other cancers.

SYLARAS software presents information on immune cell types in the form of dashboards, one per immune cell type. These dashboards summarize data on frequency, marker status, abundance in different tissue etc. as well as significant differences associated with disease progression. Dashboard attributes can be substituted with other types of information by simple changes in the SYLARAS code, making the software modular and compatible with FC, CyTOF^45^, and a set of emerging image-based single-cell analyses.^23,24,46,47,48^. We found that the scripts in SYLARAS were necessary for processing our GBM immunoprofiling data consisting of nearly 100M cells across more than 200 tissue samples. Data at this scale would be infeasible to analyze using GUI-based tools for FC data analysis such as FlowJo. SYLARAS incorporates two tools for unsupervised data clustering, PhenoGraph^17^ and FlowSOM^18^. The software also has a user interface for rapid manual gating of fluorescence histograms. These are presented in a scrolling HTML table on a Logicle scale with accompany intensity data derived from parallel control experiments. Within the SYLARAS interface it was possible to gate nearly 3,000 histograms in less than two hours without the need for automated peak finding or Gaussian mixture models.

We were surprised to find that neither FlowSOM nor PhenoGraph clustering yielded results similar to those of manual gating; in both cases, the majority of clusters contained multiple cell types and some clusters corresponded to cell types that had not previously been described. This appears to arise because unsupervised clustering is sensitive to small variations in non-specific or background fluorescence that are easily recognized as experimental artifacts by humans. In the case of high-dimensional data such as single-cell RNA-seq, unsupervised clustering is efficient, eliminates bias, and enables identification of new cell types^49^. However, in the case of antibody-based immunoprofiling, prior knowledge is already imposed at the point of antibody selection, particularly because CD antigens are often considered as dichotomous variables either present or absent on cells. Under these circumstances, gating appears to be more robust to fluctuations in background signal intensity and peak shape when compared to clustering methods, particularly when the number of samples is high and fluctuations in autofluorescence intensity accumulate. We do not doubt that a supervised workflow could be developed to recapitulate the results of human gating, but manual gating in SYLARAS is simple, reliable, and should be effective with datasets substantially larger than one presented here. We hope that the data resource on peripheral immune responses to GBM described here can serve as a foundation for more extensive characterization of systemic immunity in disease and therapy.

## ACKNOWLEDGEMENTS

This work was supported by an American Cancer Society Postdoctoral Fellowship (PF-16-197-01-LIB) to G.J.B, and by NIH/NCI grants P50-GM107618 and U54-CA225088 to P.K.S. We gratefully acknowledge G. Berriz and A. Sokolov for their computational support and S. Boswell for her preparation of the RNA-seq library.

## AUTHOR CONTRIBUTIONS

G.J.B. conceived of the project; G.J.B. and P.K.S. developed the research strategy; G.J.B., S.H.D., and J.K.M. performed the experiments; G.J.B., J.L.M., and S.K.P. wrote the source code; S.S. assisted with CyCIF and analysis of human GBM samples, G.J.B. analyzed the data; G.J.B., S.S., and P.K.S. wrote the paper.

## SUPPLEMENTAL FIGURE LEGENDS

**Supplementary Figure 1.**
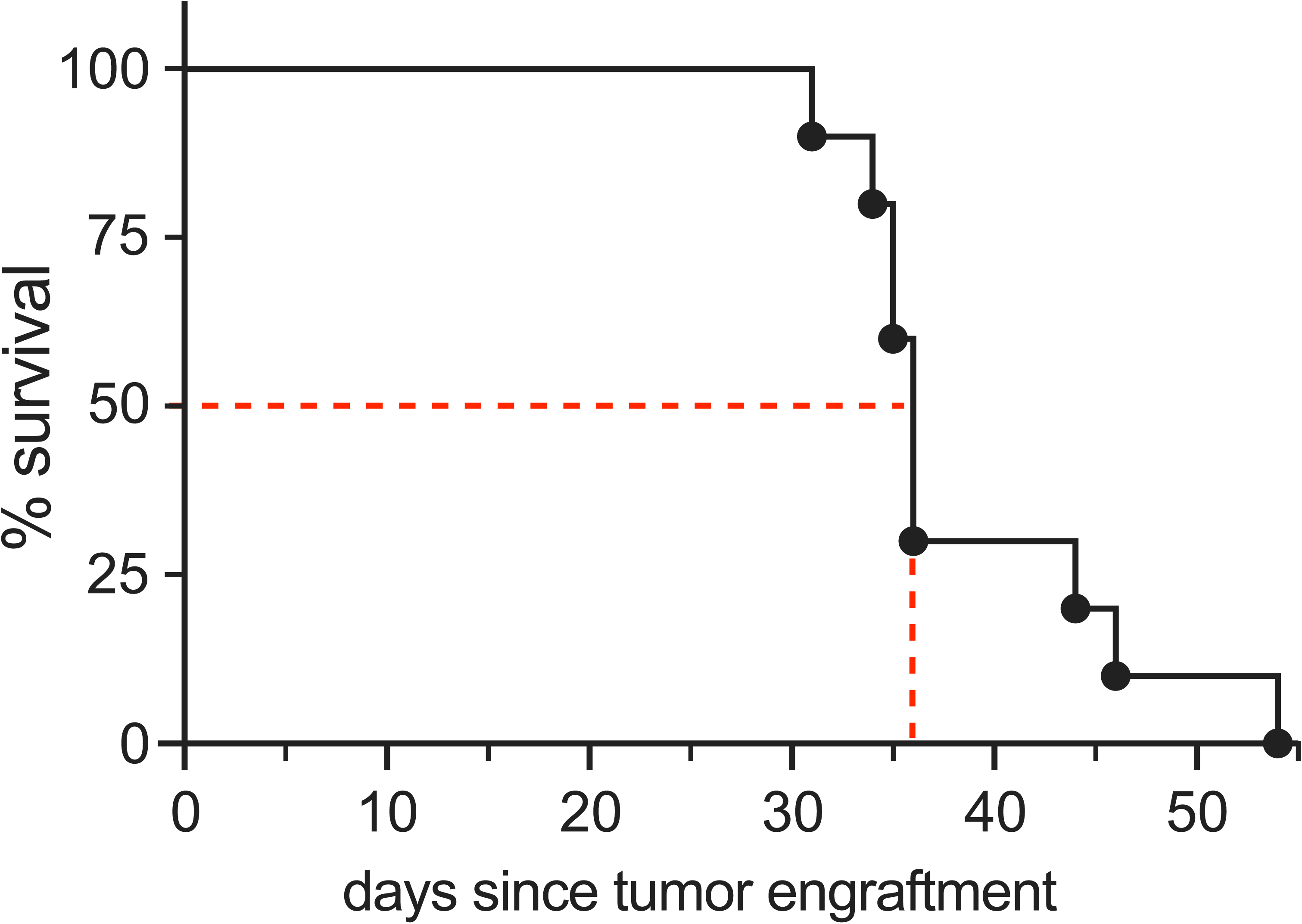
Kaplan-Meier Survival analysis of the GL261 mouse glioma model. Ten (10) C57BL/6J mice at 12-weeks-of-age were intracranially engrafted with 3×10^4^ syngeneic GL261 cells. Median survival (indicated by the intersecting dashed red lines) was 36-days post-engraftment.

**Supplementary Figure 2.**
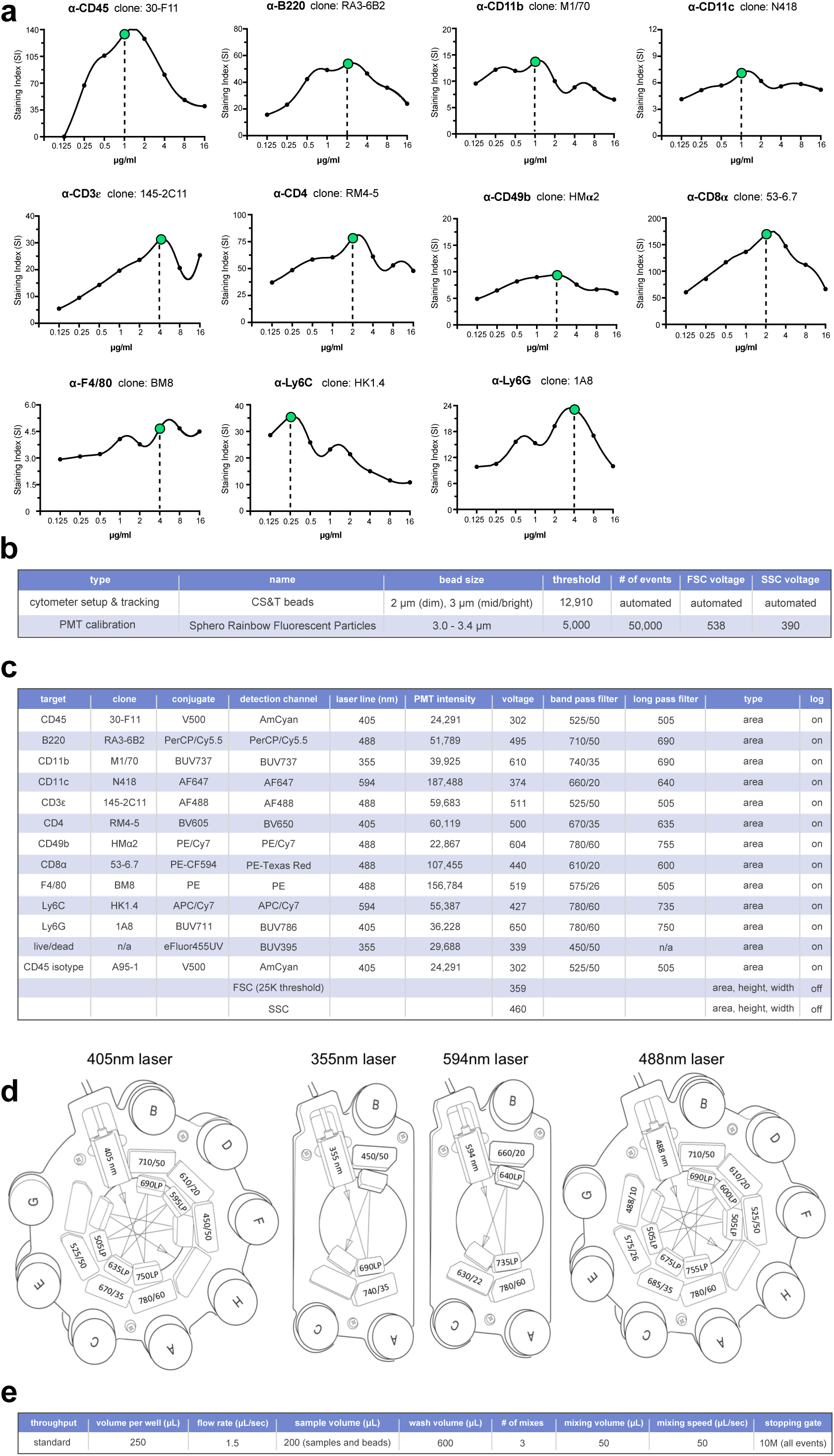
Experimental optimization for 12-color immunophenotyping by FC. (**a**) 8-point, 2-fold serial dilution titration curves for each antibody in our immunomarker panel. Splenocytes from 12-week-old female C57BL/6J mice were used in the experiment. Separation index (SI) was calculated as (MFI_pos_-MFI_neg_)/[(84%_neg_-MFI_neg_)/0.995], where MFI_pos_=median fluorescence intensity of the first positive peak, MFI_neg_=median fluorescence intensity of the autofluorescence peak, 84%_neg_=84^th^ percentile of the autofluorescence peak as previously described^50^. Cubic splines were used to interpolate adjacent data points. Antibody concentrations resulting in optimal SIs are indicated by a green dot. Data acquisition gating strategy: (FSC-A vs. SSC-A) → (SSC-H vs. SSC-W) → (FSC-H vs. FSC-W) → (DAPI-A vs. FSC-A) → (specific antibody vs. count). (**b**) Settings for cytometer setup and tracking and the photomultiplier tube (PMT) calibration used to standardize cytometer performance across three data acquisitions in this study. (**c**) Cytometer run settings. (**d**) Configuration diagrams for the BD LSR II long-pass and band-pass optical filters used in this study. (**e**) High-throughput sampler (HTS) configurations.

**Supplementary Figure 3.**
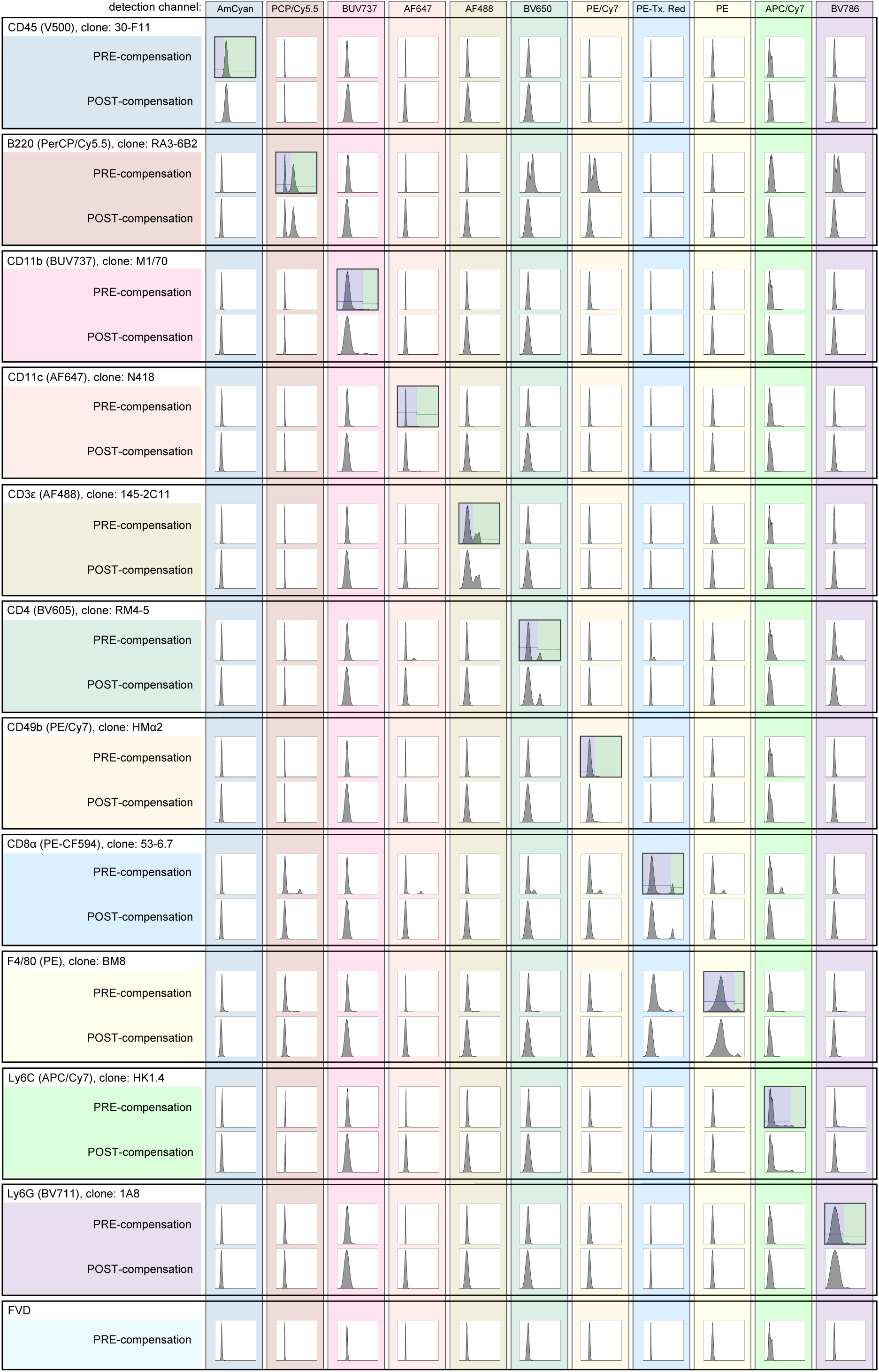
Optical spillover among an optimized panel of 12 mouse immunophenotyping antibodies is fully abrogated by spectral deconvolution. Signal intensity distributions of splenocytes from a 12-week-old female C57BL/6J mouse immunolabeled with the optimized 11 antibody immunomarker panel then stained with fixable viability dye (FVD). The 12 detection channels of a BD LSR II flow cytometer (columns) are shown pre- and post-compensation. Antibodies (rows) are color-coordinated with their target detection channel. Histograms forming the downward diagonal from left-to-right across the matrix show the respective channel’s compensation gate (blue-green interface).

**Supplementary Figure 4.**
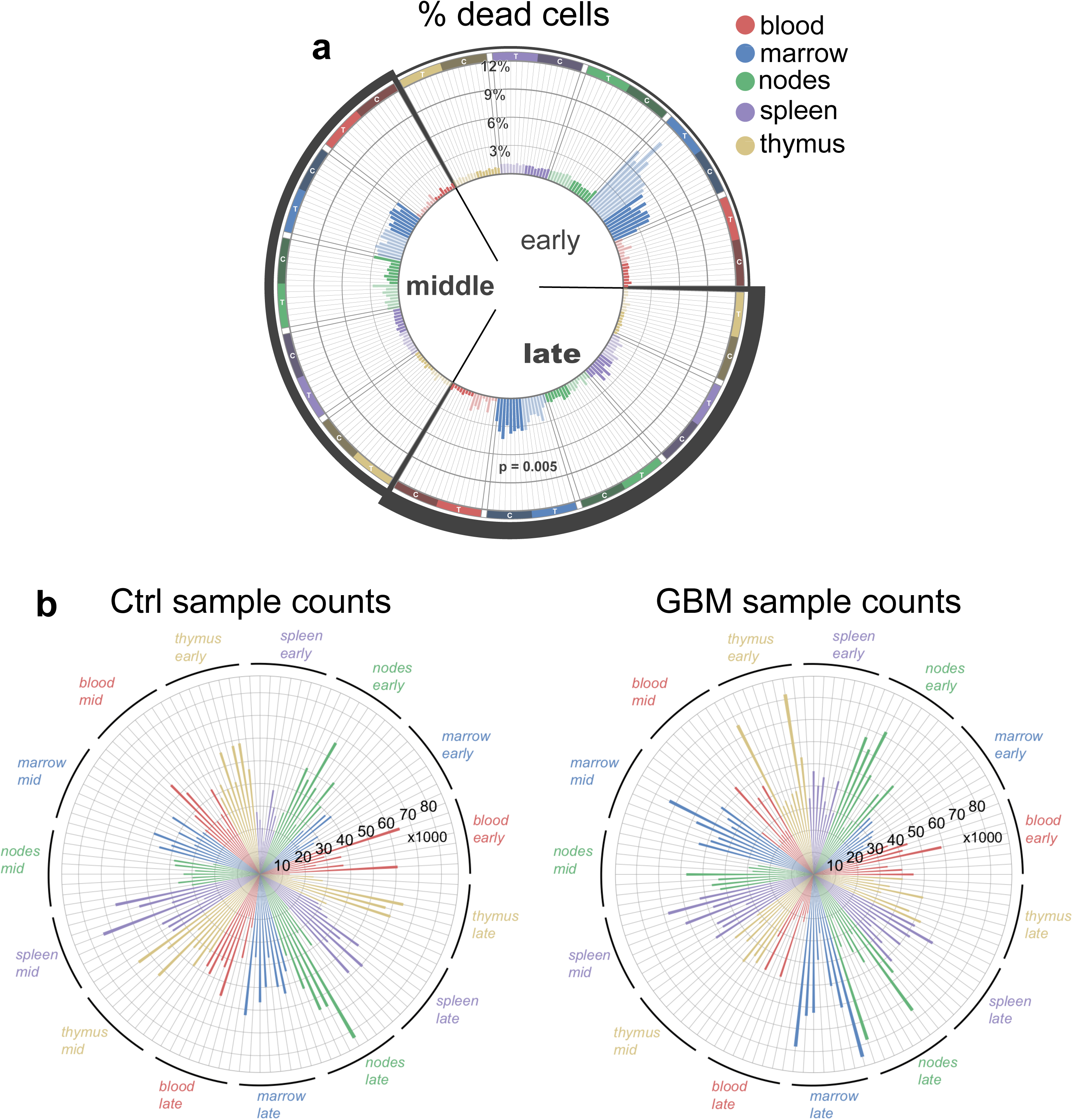
Cell viability and sample counts. (**a**) Radial bar chart showing the percentage of dead cells among the dataset’s 240 tissue samples. C=control samples, T=GBM samples. (**b**) Radial bar charts showing the number of cells in control (left) and GBM (right) tissue samples after a random sampling procedure weighted by tissue to balance the number of cells per sample.

**Supplementary Figure 5.**
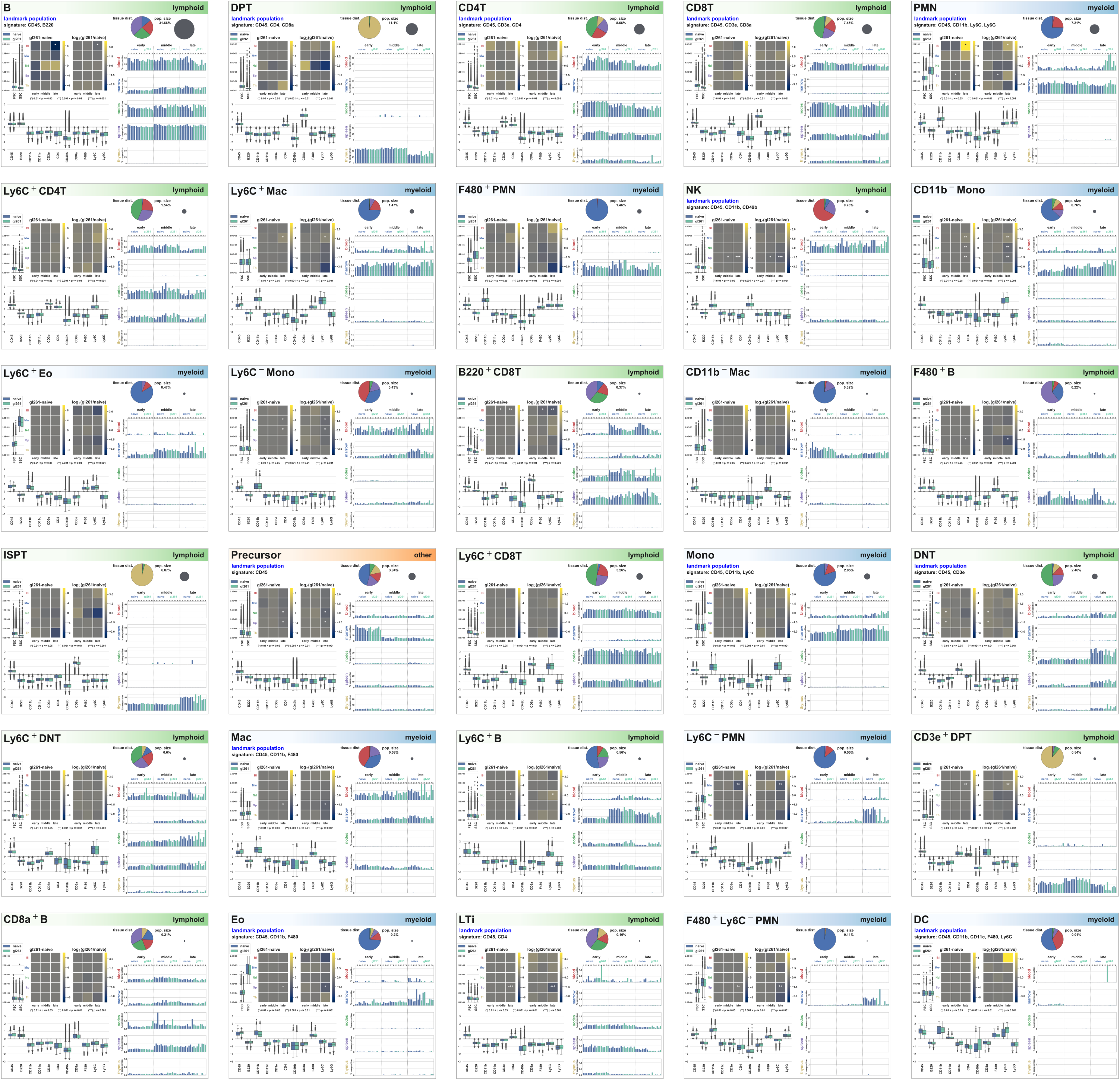
SYLARAS Dashboards. The set of 30 SYLARAS dashboards generated in this study serving as a visual compendium of time and tissue-specific perturbation of mouse immune cells in response to the GL261 mouse GBM model.

**Supplementary Figure 6.**
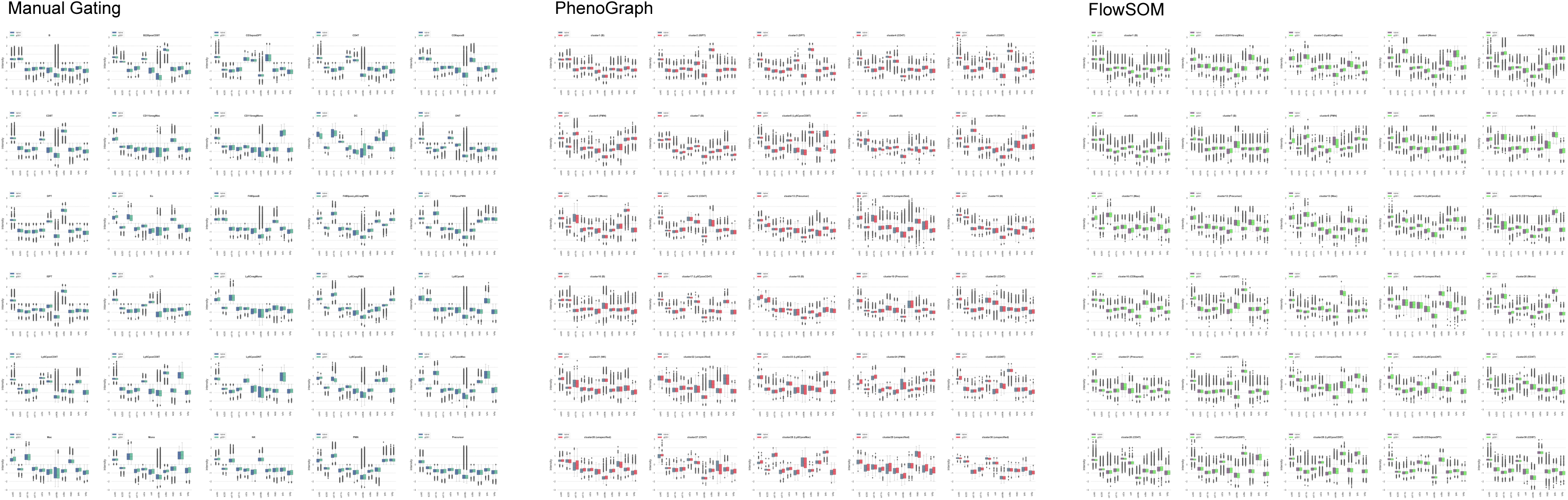
Antigen expression of different immune cell subsets identified via prior knowledge or two clustering algorithms. Boxplot distributions of Logicle-transformed antigen expression of immune cell subsets from control (naïve) and GBM-bearing (GL261) mice identified by manual gating (left), PhenoGraph clustering (middle), and FlowSOM clustering (right).

**Supplementary Figure 7.**
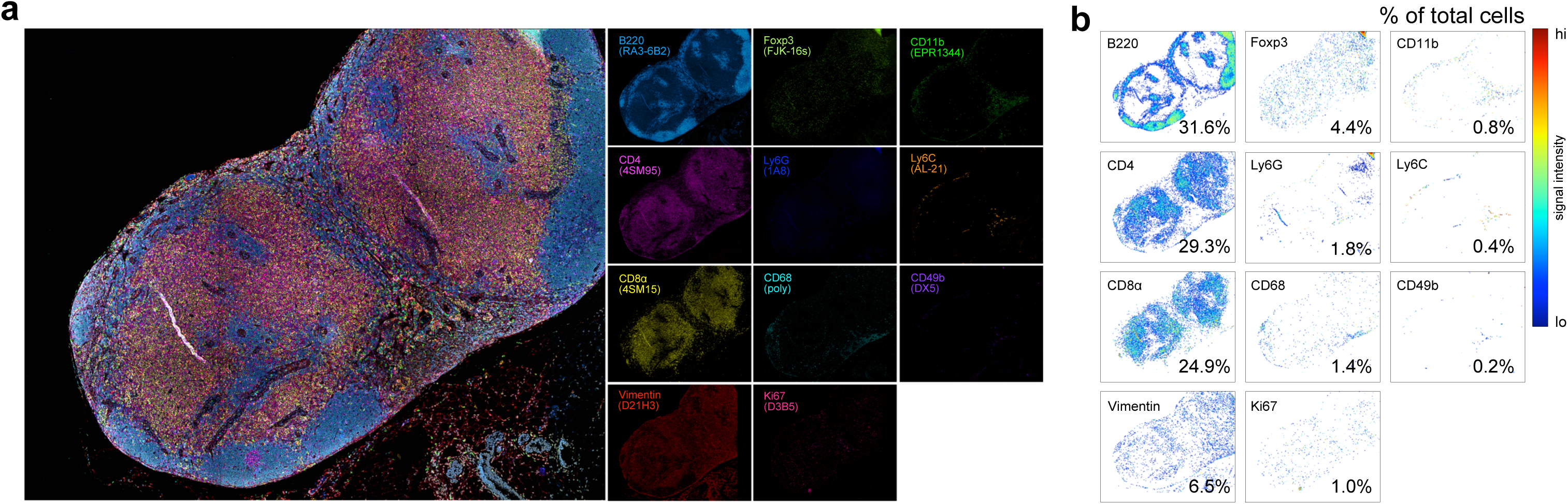
Validation of a 12-color t-CyCIF antibody panel. (**a**) 11 antibodies validated on FFPE tissue sections of a C57BL/6 mouse inguinal lymph node; composite image and individual channels are shown. (**b**) Scatter plots of single-cell data derived from the mosaic micrograph shown in panel (a) color-coded according to signal intensity of the respective antibody; the percentage of total cells immunoreactive to each antibody is shown.

**Supplementary Figure 8.**
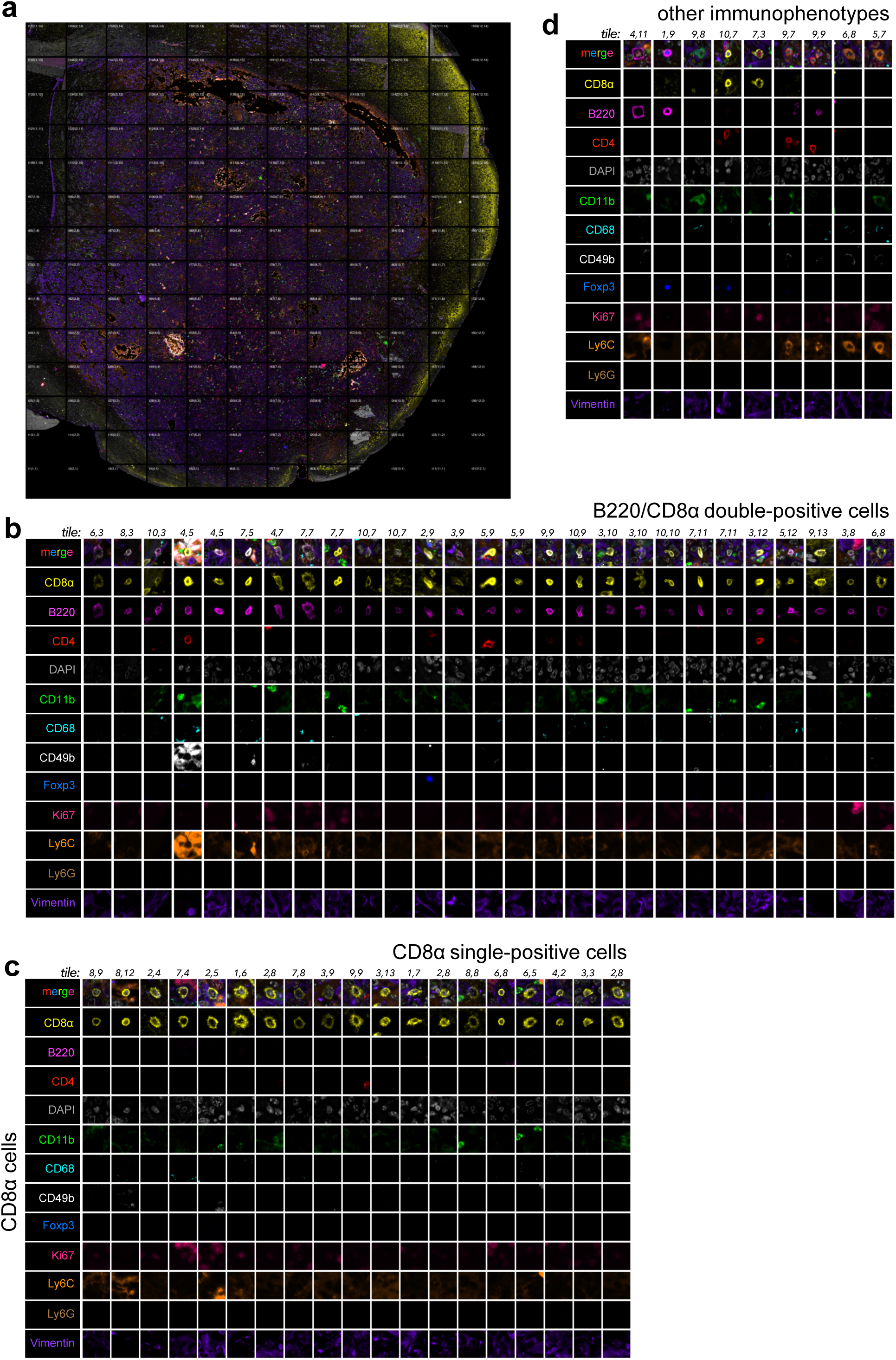
Survey of brain tumor-infiltrating lymphocytes by 12-color t-CyCIF. (**a**) A 12-channel (11 antibodies plus nuclear counter stain), 168-tile (400µm × 300µm fields of view acquired at 40x magnification) mosaic image of the tumor-ipsilateral mouse brain hemisphere bearing GL261 GBM 36-days after engraftment. Tile coordinates and number and are indicated in increasing order from the bottom-left to the top-right. Immunomarker colors are as in panels (b-d). (**b**) Examples of B220/CD8α double-positive cells. (**c**) Examples of CD8α single-positive cells. (**d**) Examples of several other low-abundance immunophenotypes identified in the GL261 TME. Tile coordinates are provided for each image for cross-referencing with the full mosaic image shown in panel (a).

**Supplementary Table 1.**
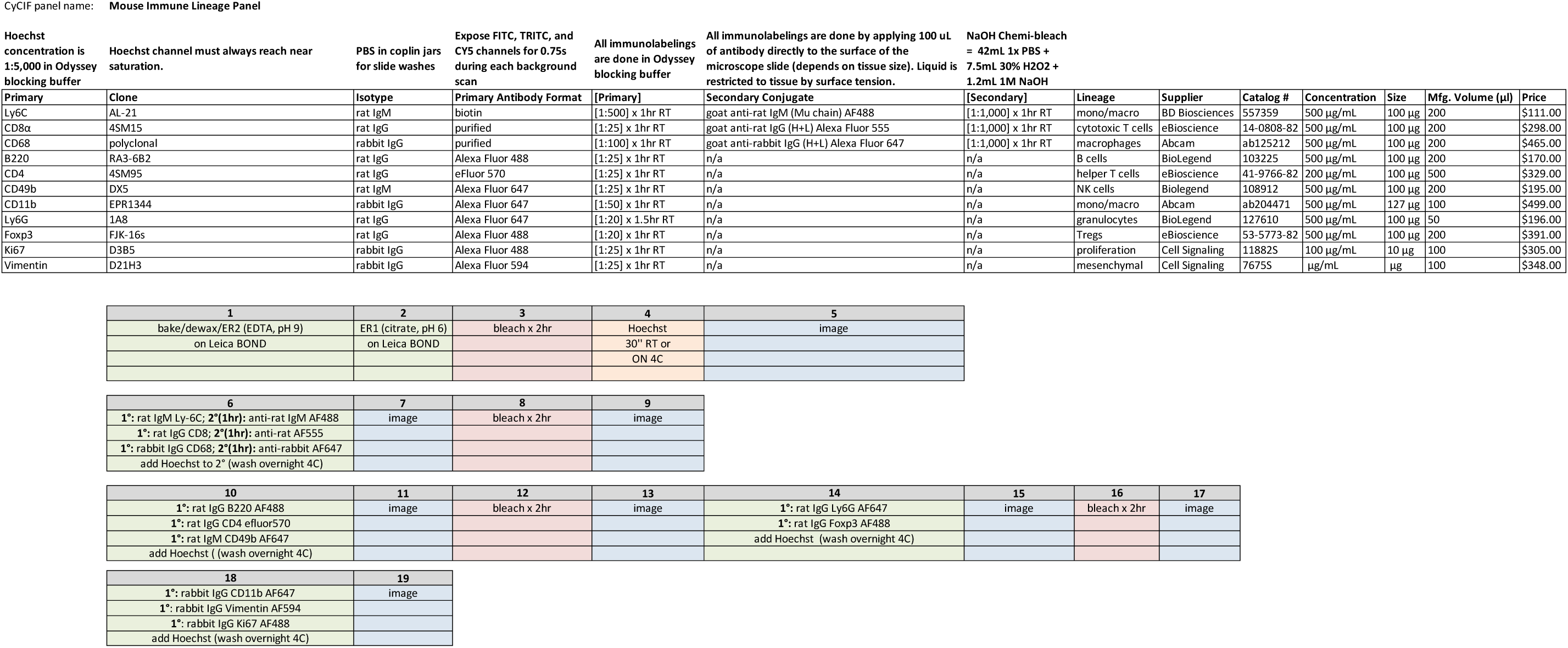
Protocol for 12-plex t-CyCIF of mouse FFPE tissue sections using immune cell lineage markers.

**Supplementary Table 2.**
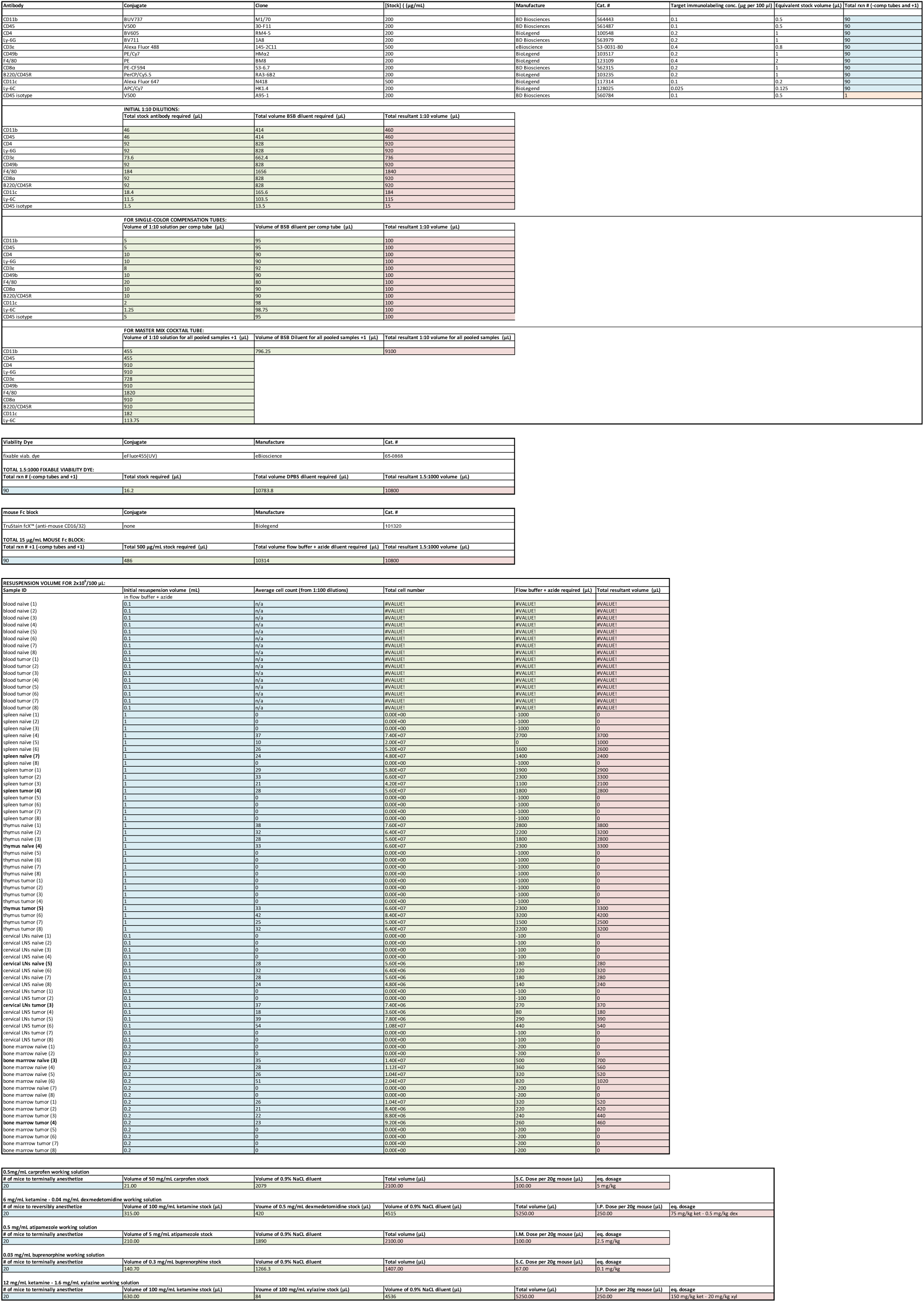
Executable spreadsheet for calculating the amounts of antibody stock and diluent, fixable viability dye, Fc block, resuspension volumes, and injectable anesthetics used in this study.

## ONLINE METHODS

Any methods, additional references, Nature Research reporting summaries, data and source code, statements of data availability, and associated accession codes are available in the online version of the paper and at https://github.com/gjbaker/sylaras and https://www.sylaras.org.

### General reagents

ddH_2_0; RPMI-1640 (Corning, Cat. No. 10-040-CV); L-glutamine (Gibco, Cat. No. 25030-081); penicillin-streptomycin (10,000 U/mL) (ThermoFisher Scientific, Cat. No. 15140-163); heat-inactivated (HI) fetal bovine serum (FBS) (Gibco, Cat. No. 16140-071); Dulbecco’s phosphate buffered saline (DPBS) w/o CaCl_2_, MgCl_2_ (Corning, Cat. No. 21-040-CV); 5% (w/v) sodium azide (NaN_3_) (BDH, Cat. No. BDH7465-2); ethylenediaminetetraacetic acid (EDTA) disodium salt dehydrate (C_10_H_14_N_2_Na_2_O_8_•2H_2_O) (Sigma Aldrich, Cat. No. ED2SS); 15 mL polypropylene conical tubes (Falcon, Cat. No. 352097); 50 mL polypropylene conical tubes (Falcon, Cat. No. 352098); 5 mL polystyrene serological pipettes (Corning, Cat. No. 4050); 10 mL polystyrene serological pipettes (Corning, Cat. No. 4100); micropipettes (1000 µL, 200 µL, 20 µL, 10 µL) (Gilson); research plus 12-channel pipette (50-300 µl), (Eppendorf, Cat. No. 3122000060); 0.1-10 µl TipOne natural pipet tips (USA Scientific, Cat. No. 1111-3200); 1.0-20 µl TipOne natural pipet tips (USA Scientific, Cat. No. 1120-1810); 1-200 µl TipOne natural pipet tips (USA Scientific, Cat. No. 1111-1200); 101-1,000 µl TipOne natural pipet tips (USA Scientific, Cat. No. 1111-2820); 40 µm nylon mesh cell strainers (Falcon, Cat. No. 352340); 2 L polyethylene Dewar flask (Nalgene, Cat. No. 4150-2000); sterile cryogenic storage vials (Sigma-Aldrich, Cat. No. V7634); mini vortexer 120V (VWR, Cat. No. 58816-121); polypropylene general-purpose test tube racks (Nalgene, Cat. No. 5930-0020); 96-well reversible microcentrifuge tube rack (Bio Plas, Cat. No. 0091); S1 pipet filler (ThermoFisher Scientific, Cat. No. 9531); 9 L TruCool rectangular ethylene-vinyl acetate foam ice pans (BioCision, Cat. No. BCS-112); 1.5 mL microcentrifuge tubes (USA Scientific, Cat. No. 1615-5500); gel loading tips (Costar, Cat. No. 4853); 60 mm × 15 mm polystyrene tissue culture dishes (Falcon, Cat. No. 353002); 0.4% trypan blue solution (Gibco, Cat. No. 15250061)

### Reagents germane to mouse euthanasia, perfusion, and tissue processing

ketamine hydrochloride injection (VEDCO, NDC: 50989-996-06); xylazine hydrochloride injection (AKORN INC, NDC: 59399-111-50); 0.9% sodium chloride (NaCl) injection, USP (Hospira, NDC 0409-4888-10); 1 mL Norm-Ject® sterile Luer-slip syringes (Henke Sass Wolf, Cat. No. 4010.200V0); PrecisionGlide needles - 26G × ½ (0.45 mm × 13 mm) (BD, Cat. No. 305111); sodium chloride (NaCl) (Sigma Aldrich, Cat. No. S9888); calcium chloride (CaCl_2_•2H_2_O) (Sigma Aldrich, Cat. No. C8106); sodium phosphate monobasic (NaH_2_PO_4_•2H_2_O) (Sigma Aldrich, Cat. No. 71505); D-glucose (C_6_H_12_O_6_) (Sigma Aldrich, Cat. No. G8270); sodium bicarbonate (NaHCO_3_) (Sigma Aldrich, Cat. No. S5761); potassium chloride (KCl) (Sigma Aldrich, Cat. No. P9333); heparin sodium salt from porcine intestinal mucosa (Sigma-Aldrich, Cat. No. H4784); extruded polystyrene foam block (2); Halsted-mosquito hemostat (2) (Fine Science Tools, Cat. No. 13008-12); fine scissors—martensitic stainless steel (2) (Fine Science Tools, Cat. No. 14094-11); Friedman rongeur (Fine Science Tools, Cat. No. 16000-14); Littauer bone cutters (Fine Science Tools, Cat. No. 16152-12); cover-glass forceps (Fine Science Tools, Cat. No. 11073-10); Dumont #5 forceps (2) (Fine Science Tools, Cat. No. 11252-40); Graefe forceps (2) (Fine Science Tools, Cat. No. 11051-10); Masterflex L/S digital pump system with easy-load II pump head, 600 RPM, 115/230V (Cole-Parmer, Cat. No. EW-77921-75); 20 G × 1 1/2” aluminum hub blunt needles (Kendall, Cat. No. 8881202363); razor blades (VWR, Cat. No. 55411-050); frosted microscope slides (Fisher Scientific, Cat. No. 12-550-343); 3 mL Luer-Lok® syringes (BD, Cat. No. 309657); PrecisionGlide needles - 23G × 1 (0.6 mm × 25 mm) (BD, Cat. No. 305145); Falcon 3 mL polyethylene transfer pipets (Corning, Cat. No. 357524)

### Reagents germane to immunolabeling

Brilliant Stain Buffer (BD Biosciences, Cat. No. 563794); TruStain FcX anti-mouse CD16/32 antibody (BioLegend, Cat. No. 101320); fixable viability dye, eFluor 455UV (eBioscience, Cat. No. 65-0868-14); Brilliant Ultraviolet 737-conjugated anti-mouse CD11b, clone: M1/70, isotype: rat DA/HA IgG2b, κ (BD Biosciences, Cat. No. 564443); V500-conjugated anti-mouse CD45, clone: 30-F11, isotype: rat LOU/M IgG2b, κ (BD Biosciences, Cat. No. 561487); Brilliant Violet 605-conjugated anti-mouse CD4, clone: RM4-5, isotype: rat IgG2a, κ (BioLegend, Cat. No. 100548); Brilliant Violet 711-conjugated anti-mouse Ly6G, clone: 1A8, isotype: rat IgG2a, κ (BioLegend, Cat. No. 127643); Alexa Fluor 488-conjugated anti-mouse CD3ε, clone: 145-2C11, isotype: Armenian hamster IgG (eBioscience, Cat. No. 53-0031-82); PE/Cy7-conjugated anti-mouse CD49b, clone: HMα2, isotype: Armenian hamster IgG (BioLegend, Cat. No. 103518); PE-conjugated anti-mouse F4/80, clone: BM8, isotype: rat IgG2a, κ (BioLegend, Cat. No. 123110); PE-CF594-conjugated anti-mouse CD8α, clone: 53-6.7, isotype: rat LOU/M IgG2a, κ (BD Biosciences, Cat. No. 562283); PerCP/Cy5.5-conjugated anti-mouse/human CD45R/B220, clone: RA3-6B2, isotype: rat IgG2a, κ (BioLegend, Cat. No. 103236); Alexa Fluor 647-conjugated anti-mouse CD11c, clone: N418, isotype: Armenian hamster IgG (BioLegend, Cat. No. 117312); APC/Cy7-conjugated anti-mouse Ly6C, clone: HK1.4, isotype: rat IgG2c, κ (BioLegend, Cat. No. 128026); V500-conjugated rat IgG2b, κ isotype control antibody, clone: A95-1, isotype: rat LOU/M IgG2b, κ (BD Biosciences, Cat. No. 560784); purified anti-human CD45R, clone: MB1, isotype: mouse IgG1 (Abnova, Cat. No. MAB16165); Alexa Fluor 488-conjugated anti-human CD8α, clone: AMC908, isotype: mouse IgG2a, κ (eBioscience, Cat. No. 53-0008-80); Alexa Fluor 555-conjugated anti-human CD3δ, clone: EP4426, isotype: rabbit IgG (abcam, Cat. No. AB208514); fixation/permeabilization solution kit (BD Biosciences, Cat. No. 554714); 4’,6-diamidino-2-phenylindole, dihydrochloride (DAPI) (ThermoFisher Scientific, Cat. No. D1306); 96-well V-bottom, non-treated, polystyrene microplate (Costar, Cat No. 3897); 12-well V-bottom reagent reservoir (Argos Technologies, Cat. No. B3135); Microseal ‘F’ foil seal (Bio-Rad; Cat. No. MSF1001)

### Reagents germane to flow cytometry

Sphero rainbow fluorescent particles (3.0-3.4 µm) (BD Biosciences, Cat. No. 556291); FACSDiva CS&T research beads (BD Biosciences, Cat. No. 655051)

### Mice

Twelve-week-old female C57BL/6J mice syngeneic to the mouse GL261 GBM model were used in this study (Jackson Laboratory, Bar Harbor, ME). All animal experiments were conducted in accordance with procedures preapproved by the Institutional Animal Care and Use Committee (IACUC) and conformed to the policies and procedures of the Center for Comparative Medicine at Harvard Medical School in agreement with the National Research Council’s “Guide for the Care and Use of Laboratory Animals”.

### GBM Cells

The GL261 GBM cell line was obtained from the Developmental Therapeutics Program (DTP), Division of Cancer Treatment and Diagnosis (DCTD) tumor repository through a material transfer agreement with the Biological Testing Branch (BTB) of the National Cancer Institute (NCI).

### Clinical Specimens

All deidentified human brain tumor resections used in this study were obtained from consented patients under care at Brigham and Women’s Hospital or Dana-Farber Cancer Institute in association with Institutional Review Board (IRB) protocols: 10-417 or 11-104 (17-000).

### Major Equipment

BD LSR II Special Order Research Product (SORP) flow cytometer w/ BD High Throughput Sampler (HTS):

- Laser line – power:

- 488 nm laser – 20 mw (run at 20 mw)
- 405 nm laser – 50 mw (run at 50 mw)
- 594 nm laser – 200 mw (run at 125 mw)
- 355 nm laser – 20 mw (run at 20 mw)

BioTek EL406 automated microplate washer/dispenser:

- In an effort to minimize cell loss, aspiration steps involving 96-well V-bottom microplates were performed using this instrument such that a 50 µL residual volume remained after each aspiration. This factor was accounted for in all reported dilutions. Instrument configurations were as follows:

- plate type: 96-well
- W-aspirate
- vacuum filtration: false
- travel rate: 1 (4.1 & 1.0 mm/sec)
- delay: 0 msec
- z-offset: 55 steps (6.99 mm above carrier)
- x-offset: 0 steps (center of well)
- y-offset: 0 steps (center of well)
- secondary aspirate : no

Beckman Coulter Avanti J-26XP centrifuge:

- JS-5.3 anodized aluminum swinging-bucket rotor (Beckman Coulter, Cat. No. 368690)

Bransonic CPXH ultrasonic cleaning bath:

- model 3800

### Reagent Preparation

supplemented RPMI-1640 (RPMI-1640 with L-glutamine, 100 U/mL penicillin-100 U/mL streptomycin, 10% HI-FBS, 0.05% sodium azide):

- to a 500 mL bottle of RPMI-1640 with L-glutamine were added:

- 5 mL of a 10,000 U/mL penicillin-10,000 µg/mL streptomycin solution
- 50 mL of HI-FBS
- 2.5 mL of a 5% w/v sodium azide solution
- 5 mL of a 10% EDTA solution

heparinized Tyrode’s solution:

- to a 1 L glass screw-cap storage bottle on a stirring plate and containing a magnetic stirring bar and 1 L ddH_2_O were added:

- 8.0 g sodium chloride
- 0.264 g calcium chloride
- 0.05 g sodium phosphate monobasic
- 1.0 g D-glucose
- 1.0 g sodium bicarbonate
- 0.2 g potassium chloride
- 100 U of heparin sodium
- Salts were allowed to completely dissolve prior to storage at 4 °C.

ammonium-chloride-potassium (ACK) lysis buffer (1X):

- to 1 L of stirring ddH_2_0 were added:

- 8.29 g of ammonium chloride
- 1.0 g of potassium bicarbonate
- 37.2 mg of sodium EDTA
- pH was adjusted to 7.4

flow buffer (DPBS + 0.5% HI-FBS):

- to 95 mL of DPBS was added:

- 0.5 mL of HI-FBS
- stored at 4°C

flow buffer + azide (flow buffer + 0.05% sodium azide):

- to 49.5 mL of flow buffer was added:

- 0.5 mL of a 5% w/v sodium azide solution
- stored at 4°C

EDTA solution (1X DPBS containing 10% EDTA):

- to 50 mL of DPBS was added:

- 5 g of disodium EDTA
- placed in ultrasonic bath to facilitate dissolution
- stored at 4°C

Fc block (flow buffer + azide containing 22.5 µg/mL anti-mouse CD16/32):

- refer to accompanying spreadsheet for preparation details (**Supplementary Table 2**).

fixable viability dye:

- eFluor 455UV fixable viability dye was diluted 1.5:1,000 in 1X DPBS
- refer to accompanying spreadsheet for preparation details (**Supplementary Table 2**).

### Antibody Titration

Immunolabeling concentrations yielding the highest staining index (SI) for each antibody in the study were determined by first harvesting the spleens of two 12-week-old female C57BL/6J mice placed under terminal anesthesia with a dose of 150 mg/kg of ketamine hydrochloride and 20 mg/kg xylazine hydrochloride diluted in sterile 0.9% NaCl delivered with a 1 mL tuberculin syringe equipped with a 26G needle as a single intraperitoneal (i.p.) injection. Using opposing frosted ends of 2 glass microscope slides, each spleen was gently macerated and rinsed with 4 mL of supplemented RPMI-1640 into a 60 × 15 mm polystyrene petri dish on ice. Splenocytes were aspirated from the dish, dispensed into a 15 mL conical tube, and centrifuged at 350 × g (max RCF) for 10 minutes at 4°C. The cell pellet was resuspended in 8 mL of a 1X ACK lysing buffer and placed on ice for 5 minutes to lyse red blood cells (RBCs). Six (6) mL of flow buffer + azide was added to the tube prior to filtering the cell suspension through a 40 µm nylon mesh into a fresh 15 mL conical tube. Splenocytes were again centrifuged at 350 × g (max RCF) for 10 minutes at 4°C then resuspended in 2 mL of flow buffer + azide. Cell counting was performed using a trypan blue and a glass hemocytometer. Cell concentration was adjusted accordingly to achieve a final concentration of 1×10^7^ cells/mL.

Two-hundred (200) µL of the 1×10^7^ cell/mL splenocyte suspension were added to wells A-H of 11 concentric columns of a 96-well V-bottom microplate using a multichannel pipette followed by centrifugstion at 100 × g (max RCF) for 3 minutes at 4°C. One-hundred and fifty (150) µL of cell supernatant was aspirated from each well followed by resuspension in 100 µL of Fc block (15 µg/mL final concentration). Splenocytes were incubated on ice for 5 minutes prior to centrifugation at 100 × g (max RCF) for 3 minutes at 4°C. One-hundred (100) µL was aspirated from each well before resuspension with 100 µL of target antibodies pre-diluted in Brilliant Stain Buffer to achieve the following two-fold serial dilution series per column for each of the 11 antibodies used in the study: 24, 12, 6, 3, 1.5, 0.75, 0.375, and 0.1875 µg/mL. Splenocytes were immunolabeled on ice for 15 minutes in the dark prior to centrifugation at 100 × g (max RCF) for 3 minutes at 4°C. Two-hundred (200) µL of supernatant were then aspirated followed by resuspension with 100 µL of flow buffer + azide. This washing step was repeated once except that 200 µL (instead of 100 µL) of flow buffer + azide was used in the resuspension step. DAPI was added to each well at a final concentration of 1µg/mL and allowed to incubate for 3-5 minutes prior to data acquisition.

Immunolabeled splenocytes were analyzed on a BD LSR II SORP flow cytometer equipped with a BD HTS for 96-well high-throughput sampling. The following acquisition gating strategy was used: (FSC-A vs. SSC-A) → (SSC-H vs. SSC-W) → (FSC-H vs. FSC-W) → (DAPI-A vs. FSC-A) → (CD_x_ vs. count). A total of 10,000 viable singlets were analyzed per well. The median fluorescence intensities (MFIs) of the first (background) and second (first true positive) peaks, and the 84^th^ percentile of the first peak were identified using the layout editor tool of FlowJo software from which a SI for each antibody was calculated according to the following formula: SI=(MFI_pos_-MFI_neg_)/[(84%_neg_-MFI_neg_)/0.995]. The maximum SI (SI_max_) of each antibody was used as the target immunolabeling concentration in our longitudinal study.

### Stereotactic Engraftment of GBM Cells into the Mouse Brain

The reader is referred to (Baker GJ et al. JoVE, 2014 PMID: 26650233) for detailed instruction on how to stereotactically engraft glioma cells into the mouse brain.

### Tissue Harvesting and Processing

The following disposable reagents were gathered and labeled before each of our study’s 3 time points:

- 15 mL conical tubes (16)

- labeled: “blood”, <condition>, <replicate> (< > indicates a variable)
- 3 mL syringes equipped with 23G needles (16)

- labeled: “marrow”, <condition>, <replicate>
- 1 mL tuberculin syringes equipped with 26G needles (17)

- (16) labeled: “blood”, <condition>, <replicate>
- (1) unlabeled: used for injectable anesthesia
- 60 × 15 mm polystyrene petri dishes (64)

- labeled: <tissue> (excluding blood), <condition>, <replicate>
- 4 mL of supplemented RPMI-1640 was added to each dish
- 0.5 mL microcentrifuge tubes (80)

- labeled: <tissue>, <condition>, <replicate>

One-hundred (100) µL of a 10% EDTA solution was added to each 15 mL conical tube. Four (4) mL of supplemented RPMI-1640 was added to each 60 × 15 mm polystyrene petri dish with the exception of those labeled “marrow”, to which only 2 mL of supplemented media was added (the other 2 mL of supplemented RPMI-1640 was to be placed into each of the (16) 3 mL syringes). One-hundred ninety-eight (198) µL of flow buffer was added to each microcentrifuge tube. Fifty (50) µL of a 10% EDTA solution was added to each 1 mL tuberculin syringe to coat the inner barrel with EDTA by operating the plunger several times. All conical tubes, petri dishes, 3 mL syringes, and microcentrifuge tubes were stored on ice or at 4°C.

Sixteen (16) mice were terminally anesthetized in series with a 150 mg/kg of ketamine hydrochloride and 20 mg/kg xylazine hydrochloride diluted in sterile 0.9% NaCl delivered with the unlabeled 1 mL tuberculin syringe as a single i.p. injection. Refer to accompanying spreadsheet for preparation details (**Supplementary Table 2**). Once non-responsive to both toe and tail pinch, each mouse was pinned ventral side up to an extruded polystyrene foam block by its front and hind paws using four 26G needles and sprayed down with 70% EtOH to prevent fur from entering the dissection cavity. Lymphoid organs were harvested in the following order: blood, thymus, spleen, superficial/deep cervical lymph nodes, bone marrow. *Blood*: after making a “y” incision from the gut to the rib cage to expose the heart, whole blood was aspirated from the right ventricle directly into one of the aforementioned EDTA-coated 1 mL tuberculin syringes. The needle was removed before expelling blood into its respectively labeled 15 mL conical tube stored on ice. Each mouse was then transcardially perfused with heparinized and oxygenated (95% O_2_/5% CO_2_) Tyrode’s solution at a rate of 4.0 mL/minute for at least 2 minutes in a laminar flow hood to fully exsanguinate the circulatory system. The method for mouse transcardial perfusion has been described in detail elsewhere (refer to citation 33 of the main text). *Thymus*: Once exsanguinated, each mouse was returned to the extruded polystyrene foam block for thymus excision with small dissection scissors and fine-tipped bent forceps being careful to remove any contaminating adipose and the mediastinal lymph nodes. Thymi were placed into respectively labeled 60 × 15 mm polystyrene petri dishes on ice. *Spleen*: Spleens were excised using small dissection scissors and fine-tipped bent forceps and placed into respectively labeled 60 × 15 mm polystyrene petri dishes on ice. *Superficial/deep cervical lymph nodes*: Lymph nodes were dissected under a dissection microscope using small dissection scissors and fine-tipped bent forceps. Excised nodes were placed into respectively labeled 60 × 15 mm polystyrene petri dishes on ice. *Bone marrow*: The right hind limb of each mouse was removed using bone cutters. Musculature and tendons were stripped away from the femur and tibia and the proximal and distal epiphyses of each bone were removed using a single-edged razor blade. Marrow from the two bones was flushed into the respectively labeled 60 × 15 mm polystyrene petri dish on ice using the 2 mL of supplemented RPMI-1640 in the respectively labeled 3 mL syringe. Flushed marrow was gently aspirated and expelled back into the petri dish one time to facilitate cell dissociation.

Thymi, spleens, and cervical lymph nodes were macerated using opposing frosted ends of 2 glass microscope slides which were dipped into the 4 mL of supplemented RPMI-1640 of the respectively labeled 60 × 15 mm polystyrene petri dish to collect as may cells as possible. Plastic Pasteur pipettes were used to transfer lymph nodes onto the frosted end of one glass microscope slide for maceration. A fresh pair of slides were used for each tissue to prevent sample cross-contamination. Five (5) mL of ice-cold DPBS was added to each petri dish prior to filtering each cell suspension through a fresh 40 µm nylon mesh into respectively labeled 15 mL conical tubes on ice using a fresh 10 mL serological pipette for every sample. Filtered samples were centrifuged at 400 × g (max RCF) for 10 minutes at 4°C. Cell supernatants were aspirated and each pellet was resuspend in 4 mL of a 1X ACK lysing buffer using a fresh 5 mL serological pipette for each sample to avoid cross-contamination. Samples were placed on ice for 5 minutes to lyse RBCs. Tissue samples were centrifuged at 400 × g (max RCF) for 10 minutes at 4°C, had their supernatants aspirated, and their cells resuspended in flow buffer + azide using the following volumes: 1000 µL for thymi; 1000 µL for spleens; 200 µL for bone marrow; 100 µL for deep/superficial cervical lymph nodes.

Blood samples were next lysed by adding 10 mL of 1X ACK lysing buffer to each 15 mL polypropylene conical tube. Tubes were placed back on ice for 5 minutes (or until blood color changed from dark burgundy to bright red). Blood samples were then centrifuged at 400 × g (max RCF) for 10 minutes at 4°C, their supernatants aspirated, and their WBC pellets resuspended with 200 µL of flow buffer + azide. Samples were stored on ice.

Two (2) µL of each tissue sample were then added to the 198 µL of flow buffer (a 1:100 dilution) in the respectively labeled 0.5 mL microcentrifuge tubes using a P20 micropipette equipped with a gel loading tip. Ten (10) µL of each 1:100 dilution was further diluted 1:1 with a 0.4% trypan blue solution. Ten (10) µL of the resultant solution was then used to estimate cell number using a brightfield microscope and a glass hemocytometer. Typical cell yields were as follows: 3×10^5^ – 1×10^6^ cells from the blood; 3×10^7^ – 8×10^7^ cells from the spleen; 6×10^7^ – 8×10^7^ from the thymus; 6×10^6^ – 1.5×10^7^ from the combined deep/superficial cervical lymph nodes; 9×10^6^ – 1.5×10^7^ from the bone marrow. Cell counts were recorded in a spreadsheet which returned the volume of additional flow buffer + azide required per sample to achieve a final concentration of approximately 2×10^7^ cells/mL (**Supplementary Table 2**). Because mouse blood typically contained less than 2×10^6^ total cells, the 200 µL volume of each blood sample was simply split between the respective experimental well and the CD49b single-positive compensation control well.

### Immunolabeling

The following disposable reagents were gathered and labeled before each of our study’s 3 time points:

- 1.5 mL microcentrifuge tubes (12)

- labeled in duplicate: <target antibody or CD45 isotype>
- tubes were placed into a microcentrifuge tube rack on ice
- 12-well V-bottom reagent reservoir (1)

- individual compartments labeled: <target antibody or CD45 isotype>
- 15 mL conical tubes (2)
- one labeled “cocktail”, the other labeled “FVD”
- placed on ice

Dilutions of each antibody (1:10) were prepared using the 12 microcentrifuge tubes. Refer to accompanying spreadsheet for preparation details (**Supplementary Table 2**). From these 1:10 dilutions, antibodies were further diluted in the respective wells of a 12-well V-bottom reagent reservoir to achieve 100 µL volumes at the final antibody concentration to immunolabel the cells of each single-color compensation control. The remaining 1:10 antibody dilutions were used to prepare a master mix of combined target antibodies (excluding the CD45 isotype) placed in the 15 mL tube labeled “cocktail”. All final antibody dilutions were stored on ice in the dark.

One-hundred (100) µL aliquots of each cell suspension were added to the wells of a 96-well V-bottom microplate using a multichannel pipette according to the plate diagram shown in (**Fig. 1**). The plate was then centrifuged at 100 × g (max RCF) for 6 minutes at 4°C. Fifty (50) µL of supernatant was aspirated and cells were resuspended with 100 µL of Fc block (15 µg/mL final concentration) the total volume of which was determined using the spreadsheet (**Supplementary Table 2**). Cells were blocked on ice for 5 minutes before being centrifuged at 100 × g (max RCF) for 5 minutes at 4°C. Fifty (50) µL of cell supernatant was aspirated. The 80 experimental tissue samples were resuspended with 91 µL of the combined target antibodies from the tube labeled “cocktail”. The 100 µL volumes in the 12-well V-bottom reagent reservoir were added to the respective single-color compensation control wells; CD45 isotype antibodies were added to the “ISO” well. Cells in the “UNS” and “FVD” wells were resuspended with 91 µL of Brilliant Stain Buffer. The microplate was allowed to incubate on ice in the dark for 15 minutes before adding 100 µL of DPBS to each well and mixing thoroughly with a multichannel pipette. The plate was then centrifuged at 100 × g (max RCF) for 5 minutes at 4°C followed by aspirating 191 µL from each well. Two-hundred (200) µL of DPBS were added to each well using a multichannel pipette and mixed thoroughly by pipetting. The plate was centrifuged at 100 × g (max RCF) for 5 minutes at 4°C followed by a 200 µL aspiration of supernatant. Fixable viability dye was diluted 1.5:1,000 in 1X DPBS in the 15 mL conical tube labeled “FVD” according to the spreadsheet (**Supplementary Table 2**). One-hundred (100) µL of the diluted FVD were added to each well (a 1:1,000 final dilution) except for the “UNS”, which was resuspended with 100 µL of DPBS only. Cells were incubated on ice in the dark for 30 minutes. One-hundred (100) µL of DPBS were then added to each well and with a multichannel pipette and pipetted thoroughly to wash. The plate was centrifuged at 100 × g (max RCF) for 5 minutes at 4°C followed by a 200 µL aspiration, addition of 200 µL of DPBS using a multichannel pipette, and through mixing. The plate was centrifuged at 100 × g (max RCF) for 5 minutes at 4°C followed by a 200 µL aspiration.

One-hundred (100) µL of a fixation/permeabilization solution were next added to each well and immediately resuspended with a multichannel pipette to prevent cell-to-cell crosslinking. Cells were incubated on ice in the dark for 20 minutes followed by the addition of 100 µL of flow buffer and thorough mixing with a multichannel pipette. The plate was centrifuged at 100 × g (max RCF) for 5 minutes at 4°C followed by a 200 µL aspiration and resuspension with 200 µL of flow buffer. A Microseal ‘F’ foil seal was applied to the top of the 96-well microplate to prevent dehydration, wrapped in aluminum foil to block light, and stored at 4°C prior to data acquisition by flow cytometry.

### PMT Calibration

Detection channel PMT voltages were calibrated such that the signal intensity distribution corresponding to background autofluorescence of FVD-labeled splenocytes was on scale and to the left of center. FVD-labeled single-color compensation control splenocytes were run to verify that the assigned PMT voltages were compatible with the dynamic range of each immunomarker’s expression profile (i.e. that immunopositive cells were on scale). To prevent downstream compensation values from exceeding 100%, optical spillover of each single-color compensation control into off-target detection channels was checked to ensure that peak signal intensity occurred in the target detection channel. Sphero Rainbow Fluorescent Particles (single-positive beads) were run to predefine tolerability ranges for laser intensity, stability, and alignment before initiating our study so that changes in laser emission power could be monitored and accounted for between successive data acquisitions to prevent run-to-run variation. Single-positive beads were gated according to the following strategy:

- FSC-A vs. SSC-A: on single-positive beads
- biexponential histograms of all detection channels
- An interval gate was placed around the peak in each detection channel to define a narrow tolerability range to calibrate to between runs.

### Data Acquisition

Cytometer setup & tracking was performed using FACSDiva CS&T research beads to optimize and standardize instrument performance prior to each data acquisition. The 96-well V-bottom plate containing the 80 immunolabeled experimental tissue samples and 16 optical controls was then loaded into a BD HTS affixed to a BD LSR II SORP flow cytometer. The gating strategy used at each acquisition was as follows:

- FSC-A vs. SSC-A: on all events (minus RBCs/debris)
- SSC-H vs. SSC-W: doublet discriminator
- FSC-H vs. FSC-W: doublet discriminator
- BUV395-A (FVD) (detected with 355 nm laser off of a 450/50 band pass filter) vs. FSC-A: on FVD negative cells (i.e. viable cells)
- biexponential histograms of all detection channels

- laser line, band pass filter, long pass filter, antibody detected:

- 405 nm, 525/50, 505, V500-CD45
- 488 nm, 710/50, 690, PerCP/Cy5.5-CD45R/B220
- 355 nm, 740/35, 690, BUV737-CD11b
- 594 nm, 660/20, 640, Alexa Fluor 647-CD11c
- 488 nm, 525/50, 505, Alexa Fluor 488-CD3ε
- 405 nm, 670/35, 635, BV605-CD4
- 488 nm, 780/60, 755, PE/Cy7-CD49b
- 488 nm, 610/20, 600, PE-CF594-CD8α
- 488 nm, 575/26, 505, PE-F4/80
- 594 nm, 780/60, 735, APC/Cy7-Ly6C
- 405 nm, 780/60, 750, BV711-Ly6G

Samples were run in the following order:

1. SP beads (PRE): to check that PMT voltages were within previously defined tolerability ranges prior to data acquisition.
2. unstained control splenocytes (UNS)
3. unstained control splenocytes labeled with FVD (FVD)
4. control splenocytes labeled with FVD and CD45 isotype control antibodies (ISO)
5. single-color compensation controls stained with FVD (control splenocytes were used for each antibody except CD49b, in which case WBCs were used due to the increased abundance of CD49b^+^ cells in blood)
6. experimental samples (16 mice × 5 tissue samples=80 total)
7. SP beads (POST): to check that PMT voltages remained stable over the acquisition period (fluidic anomalies can impact laser delay stability)

Raw data were exported as FCS3.0 files upon completion of each data acquisition session.

### Spectral Deconvolution and Data Preprocessing

FlowJo software was used to spectrally deconvolve raw flow cytometry data. Data from 15 of the 16 optical control wells (UNS was excluded) were imported into the *compensation group* of a new FlowJo *workspace*. CD45 signal intensity distributions were invariably unimodal making it difficult to objectively define a compensation gate for the CD45 single-positive control alone. Thus, the data corresponding to well E11 (ISO) was merged with that of F9 (CD45 single-color compensation control) using FlowJo’s *concatenate* feature. The merged data were saved as a new FCS3.0 file and imported into the *compensation group* of the current *workspace*. The original CD45 single-positive compensation control and ISO samples were then deleted from the *workspace*. The merging procedure resulted in a bimodal distribution and the ability to objectively define a CD45 compensation gate between its two peaks. The merged CD45 file plus the other 10 single-color compensation controls and the FVD well (E10)—which served as the compensation control for the FVD itself—were gated for viable singlets according to the following strategy:

- FSC-A vs. SSC-A: on all events (minus RBCs/debris)
- SSC-H vs. SSC-W: doublet discriminator
- FSC-H vs. FSC-W: doublet discriminator
- BUV395-A (FVD) vs. FSC-A (viewed as a contour plot at the 2% level): on FVD negative cells (i.e. viable cells)
- backgate to FSC-A vs. SSC-A: on total viable singlets (or a subset for scarce populations)

Once gated, viable singlets of each compensation control sample were visualized as histograms in their respective detection channel. Signal intensity values of each histogram were bisected at the interface of the penultimate and ultimate modes of each signal intensity distribution using FlowJo’s *bisector tool*. Its *compensation* tool was then opened and the subsets to the left and right of the bisection were dragged into fields labeled *negative* and positive, respectively. The process was repeated for all 11 antibodies plus the FVD. Next, a new *group* in the *workspace* window was generated and titled “cocktails” to which the 80 experimental samples were imported. The finalized compensation matrix was then applied to the “cocktails” group. Viable singlets from each experimental sample were gated in the same way as the single-color compensation controls according to the first 4 steps of the gating strategy outlined above then exported as new FCS3.0 files; the aggregate of such files from each of the study’s 3 time points served as input into our computational data analysis algorithm.

### Weighted Random Sampling

A 10 million cell weighted random sample (WRS) was derived from the preprocessed flow cytometry data to help balance the number of cells per tissue sample. Sample weights were defined per tissue per cell according to the formula 1/(ω × N*_i_*) where ω is the number of unique tissue types (5 in this cases) and N*_i_* is the number of events associated with the *i*^th^ tissue (blood, marrow, nodes, spleen, and thymus).

### Histogram Gating

Preprocessed flow cytometry data (i.e. compensated viable singlets) were displayed as 2,640 histograms (240 samples × 11 antibody detection channels) plotted on a Logicle scale and formatted as scalable vector graphics (SVGs) in the context of a scrolling HTML table viewable with a web browser. A KDE of the signal intensity distribution of cells from the FVD well (i.e. compensated unstained viable splenocytes) was superimposed on each to identify signal intensity values corresponding to autofluorescence background. This allowed for the rapid recording of gate values at the interface of the first (autofluorescence) and second (true signal) peaks for all histograms in a .TXT file. A vertical line was then programmatically rendered at the location of each gate value and again visualized as a scrolling HTML table on the web for confirmation or refinement. The numerical value of each gate value was then Logicle-transformed and subtracted from the Logicle-transformed signal intensity values of its corresponding histogram (i.e. Logicle[signal intensity value_i_]) – Logicle[gate value]) resulting in the Logicle-transformed gate value assuming the numerical value of zero and background signal intensity values becoming negative valued. Since the 5 lymphoid tissue types predominately consisted of immune cells, the CD45 signal intensity distributions were invariable unimodal with no discernable local minima. Thus, for each time point and tissue combination, a common CD45 bias was curated by pooling the corresponding samples, computing Q25 – [1.5 * [Q75 - Q25]], then rounding to the nearest multiple of 5. (where Q25 and Q75 were the first and third quartiles of the Logicle-transformed data, respectively).

Immunophenotypes interpreted as CD49b^+^ granulocytes were identified in the blood of both control and GBM mice. Since granulocytes interact with CD49b^+^ platelets, we considered these cells as a likely artifact of residual platelets present within blood samples. In taking a conservative approach to correct for this discrepancy, cell status of the CD49b immunomarkers was only considered in cases where the immunophenotype was otherwise consistent with NK cells (e.g. CD45^+^, CD49b^+^, CD11b^+^). Thus, a Boolean truth value of zero for the CD49b channel was uniformly applied to all cells whose immunophenotype was not matching NK cells.

### t-CyCIF

The reader is referred to www.cycif.org for details on the t-CyCIF methodology. Briefly, 5µm-thick FFPE tissue sections of the tumor-ipsilateral brain hemisphere of a C57BL/6J mouse engrafted with 3×10^4^ GL261 cells 36-days prior was serially scanned using 2×2 binning with a CyteFinder slide scanning fluorescence microscope (RareCyte, Seattle, WA, USA) equipped with a 40X (0.6NA) long working distance objective on each of 4 imaging cycles. Before immunolabeling with the first set of 3 fluorophore-conjugated antibodies, the tissue was counterstained with DAPI at a 1:5,000 dilution of a 10 mg/mL stock in 1x PBS for 30 minutes at RT then blocked with Odyssey Blocking Buffer at RT for 1 hour to limit non-specific antibody binding. Blocked slides were then imaged to document background autofluorescence prior to immunolabeling. Antibody incubations were performed at 4°C overnight in an opaque and humidified chamber. After immunolabeling, slides were washed 3 times with 1x PBS, temporarily coverslipped in 1x PBS containing 10% glycerol, and re-imaged. Immediately after imaging, slides were de-coverslipped by vertical submersion in Coplin jars containing 1x PBS until the coverslip spontaneously fell way from the slide. Antibody fluorescence was deactivated by submersion of the slide in a 3% H_2_O_2_ solution containing 20 mM NaOH in 1x PBS for 2 hours at RT in the presence of intense fluorescent light. Following fluorophore deactivation, slides were rinsed briefly in 1x PBS, then subject to the next round of immunolabeling.

Upon completion of the cyclic imaging procedure, autofluorescence background was computationally subtracted on a pixel-by-pixel basis using ImageJ. The final 12-plex mosaic image was generated by aligning (or registering) the 168 400×300µm imaging fields acquired during each imaging cycle on their DAPI counterstained nuclei using ImageJ’s Multistack Registration Plugin. Registered images were compiled into multi-image stacks and segmented based on the DAPI signal acquired during the last imaging cycle by applying a uniform threshold on the DAPI channel across all images and then converting this signal into binary regions of interest (ROIs) using the ImageJ’s *Analyze Particles* function. ROIs corresponding to single-cell nuclei were enlarged by 3 pixels to cover immune cell cytoplasm and membrane. Respective ROIs where overlaid on each image to obtain integrated fluorescence signal intensity data on each cell for across all 12 immunofluorescence channels.

### RNA-seq

The spleens of five 12-week-old tumor-naïve C57BL/6J mice were immunodepleted of B cells using MojoSort mouse CD19 nanobeads (Biolegend, Cat. No. 480002) before FACS-purifying B220^+^ CD3ε^+^ CD8α^+^ (i.e. B220^+^ CD8T cells) and B220^−^ CD3ε^+^ CD8α^+^ (i.e. CD8T cells) into respective 15 ml conical tubes containing ice-cold with 0.5% BSA DPBS. Approximately 1×10^6^ cells of each type were centrifuged at 400 × g (max RCF) for 10 minutes at 4°C and resuspended in RLT lysis buffer prior to total RNA extraction with Qiagen’s RNeasy mini kit using the optional DNase treatment (Qiagen, Cat. No. 74104). Total RNA was then used to prepare sequencing libraries of the coding transcriptome using Illumina’s TruSeq stranded mRNA protocol. Libraries were sequenced by synthesis on a Nextseq 500 instrument. Fastq files were processed on a high-performance computer cluster using a standardized data analysis pipeline involving the analysis programs FASTQC, STAR, Salmon, featureCounts, EdgeR, Kallisto. Kallisto transcript abundance files were then analyzed on a desktop computer using the Sleuth RNA-seq analysis program.

### Software

- FACSDiva (version 8.0)
- FlowJo (version 10.3.0)
- Python (version 3.6.1)

### Statistics

Statistical tests were performed using SciPy.stats, a Python-based library of validated statistical functions. Statistical tests used throughout this paper are described at their point of reference. Two-sample hypothesis tests were considered statistically significant if their FDR-adjusted p-value (q-value) reached a cutoff of ≤0.05.

### Data and Code Availability

SYLARAS source code and the GBM flow cytometry dataset are freely available for download at GitHub (https://github.com/gjbaker/sylaras), Synapse (https://www.synapse.org/#!Synapse:syn21038618), and the SYLARAS project website (https://www.sylaras.org).

